## Supplementary File I for "STANCE: a unified statistical model to detect cell-type-specific spatially variable genes in spatial transcriptomics"

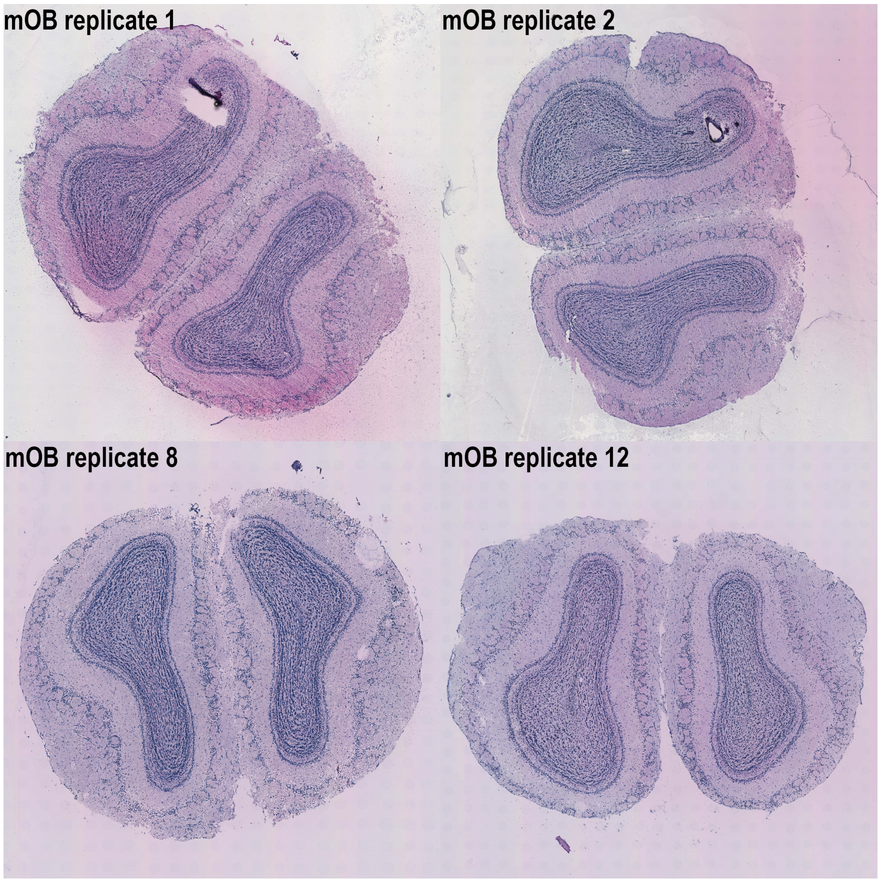


Figure S1: **Images of four mouse olfactory bulb tissue sections.** Displayed are the Hematoxylin and eosin-stained brightfield images of mouse olfactory bulb tissue replicate sample 1, 2, 8, and 12[1] (https://www.spatialresearch.org). The variability in tissue orientation and positioning during sample preparation highlights the need for rotation-invariant statistical methods in spatial transcriptomics.

### Spatial Rotation Simulation

We did a simulation to demonstrate that CSIDE, spVC and CTSV have statistical issues in ctSVG detection, due to the fact that they treat spatial locations as fixed effect and such analysis are not spatial rotation-invariant.

#### Import paksges and the sample pattern

```
library(ggplot2)
library(dplyr)
library(tidyr)
library(spacexr)
library(CTSV)
library(spVC)
library(SpatialExperiment)
library(Triangulation)
```

#### Simulate single cell resolution spatial transcriptomics data

We first simulated single cell resolution spatial transcriptomics data. We borrowed the tissue shape of the mouse olfactory bulb (MOB) dataset [1] and generated 3000 single cells as well as 3 spatial domains through the `SRTsim`[2] package. (Here, the pre-simulated single cell data `Sample.rda` is available at https://drive.google.com/drive/folders/1KSxeInbwFswuJdTBxCjMZKc6voz5UWqZ?usp=drive_link.)

```
set.seed(1)
load(file = "./Sample.rda")
# number of single cells
numCells <- nrow(dat.sc)

plot_spatial_pattern <- ggplot(dat.sc, aes(x, y, color = domain)) +
  geom_point() +
  theme_minimal() +
  theme(legend.title = element_text(size = 11, face = "bold"),
        panel.grid.major = element_blank(),
        panel.grid.minor = element_blank()) +
  labs(color = "Domain") +
  guides(color = guide_legend(override.aes = list(size = 4)))

print(plot_spatial_pattern)
```

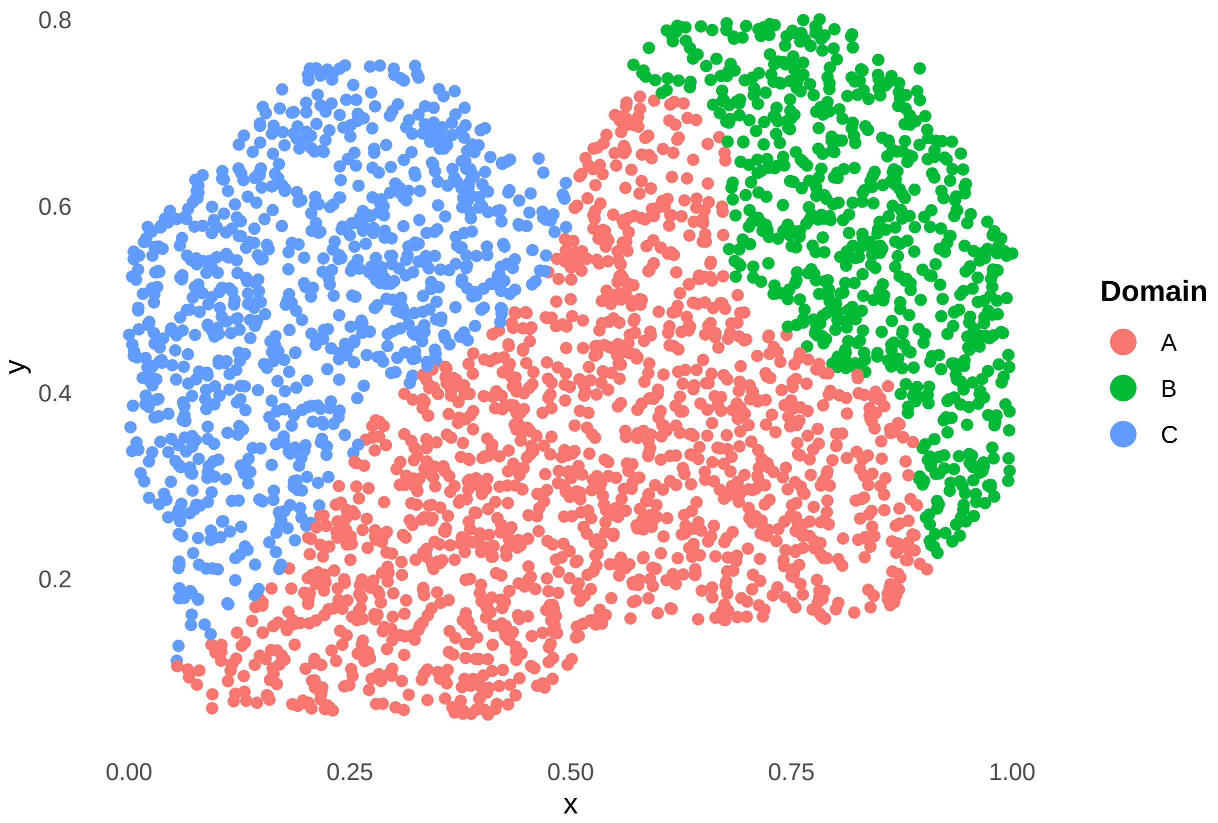


Each cell is assigned to one of three cell types based on a categorical distribution, with probabilities of 40% for cell type 1 (CT1), 30% for cell type 2 (CT2), and 30% for cell type 3 (CT3).

```
# Assign single cells into 3 cell type groups
Cell_Types <- paste0("CT",extraDistr::rcat(n = numCells,
                                           prob = c(0.4, 0.3, 0.3)))
dat.sc$cell_type <- factor(Cell_Types)
plot_CT_pattern <- ggplot(dat.sc, aes(x, y, color = cell_type)) +
  geom_point() +
  theme_minimal() +
  theme(legend.title = element_text(size = 11, face = "bold"),
        panel.grid.major = element_blank(),
        panel.grid.minor = element_blank()) +
  labs(color = "Cell types") +
  guides(color = guide_legend(override.aes = list(size = 4)))
print(plot_CT_pattern)
```

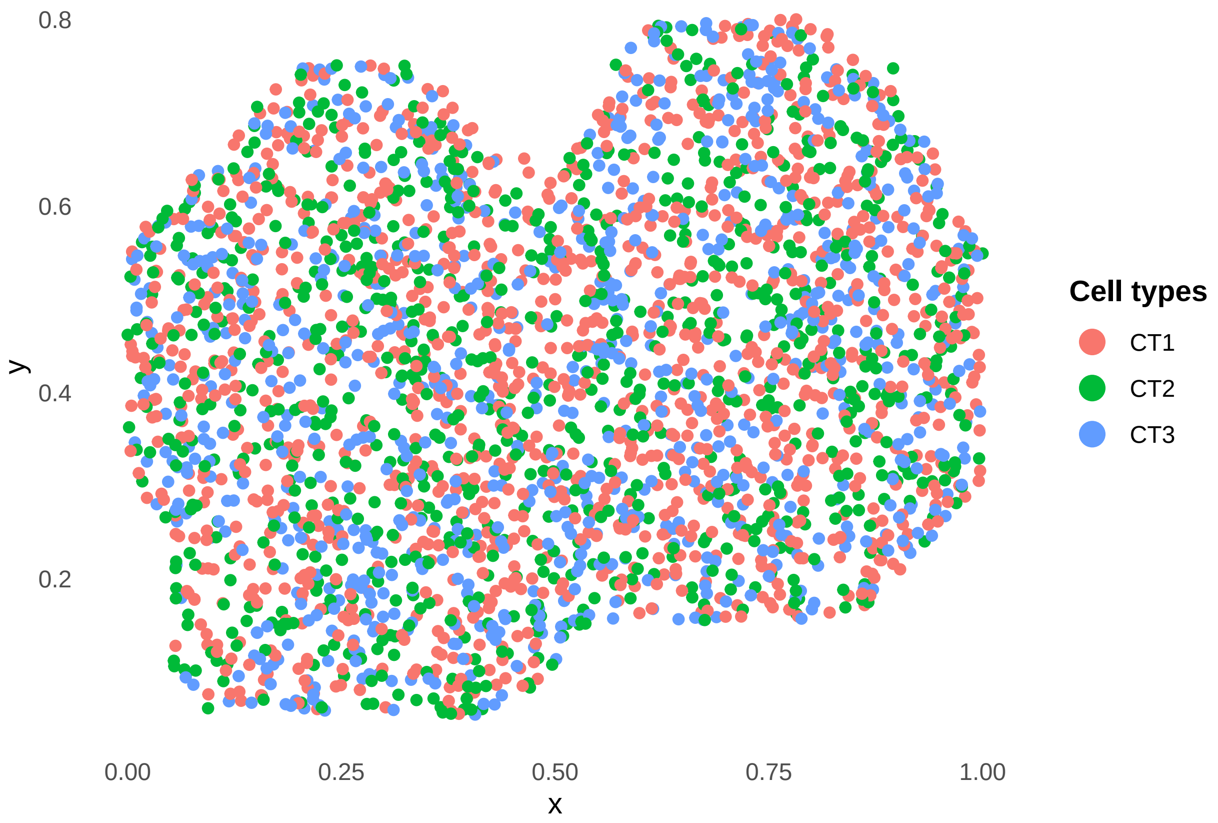


We assumed the presence of 3 distinct cell types and simulated the expression of 50 genes per cell using a negative binomial distributions characterized by mean 1 and dispersion parameter 1.5.

```
# number of genes
numGenes <- 50
# mean and dispersion parameter of negative binomial distribution
mu <- 1
dispersion <- 0.7
# Baseline expression
counts.null <- matrix(rnbinom(n = (numGenes * numCells),
                              size = dispersion,
                              mu = mu),nrow = numGenes)
counts.sc <- counts.null
```

We selected 30 out of 50 genes serving as cell type marker genes, in which each of three cell types has 10 unique marker genes. For each specific cell type, we modified the expression of their marker genes with a fold change of 4 (i.e., multiplying the mean parameter of the negative binomial distribution by 4), regardless of their domain assignments.

```
# Fold change the mean by 4 to the expression of 10 marker genes for each cell type
for (iCT in 1:3){
  counts.upregulated <- matrix(rnbinom(n = (10 * numCells),
                                       size = dispersion,
                                       mu = mu * 4),nrow = 10)
  cell_fold_change.1 <- which((dat.sc$cell_type == paste0("CT",iCT)))
```

```
  counts.sc[(iCT*10 - 9):(iCT*10),
            cell_fold_change.1] <- counts.upregulated[,cell_fold_change.1]
}
```

Next, we selected another 10 genes, Gene 31 to Gene 40, to serve as CT1-specific ctSVGs. For cell type CT1, we adjust the mean expression of the genes with a fold change of 4 for cells located outside domain $A$.

```
cell_fold_change.2 <- which((dat.sc$domain != "A") & (dat.sc$cell_type == "CT1"))
# Fold change the mean by 4 to the expression of 10 SVGs
# for the single cells within domain B & C
counts.sc[31:40,cell_fold_change.2] <- matrix(
  rnbinom(n = (10 * length(cell_fold_change.2)),size = dispersion,mu = mu * 4),
  nrow = 10)

row.names(counts.sc) <- paste0("Gene", 1:dim(counts.sc)[1])
colnames(counts.sc) <- paste0("Cell", 1:dim(counts.sc)[2])
```

The tissue was divided into 251 spots using a grid size of 0.05. For each spot, we aggregated the expression counts of all cells within it to determine spot-level expression and calculated cell type compositions. Additionally, the coordinates of each spot were based on the mean x and y coordinates of the cells within that spot.

```
## Aggregate to obtain spot-level gene expression
counts.st <- apply(counts.sc, MARGIN = 1,
                   FUN = function(gene_expr, dat = dat.sc, grid.size = 0.05){
                     # Combine gene expression vector into dat
                     dat <- cbind(dat, gene_expr)

                     # Calculate grid indices for each cell
                     dat$grid_x <- floor(dat$x / grid.size)
                     dat$grid_y <- floor(dat$y / grid.size)

                     # Aggregate gene expression by grid and calculate center of mass
                     expression_summary <- dat %>%
                       group_by(grid_x, grid_y) %>%
                       summarise(
                         expr = sum(gene_expr),
                         .groups = 'drop'
                       )
                     return(expression_summary$expr)
                   })
counts.st <- t(counts.st)

# number of spots
n <- ncol(counts.st)

# Calculate proportions of each cell type within each grid
cell_type_counts <- dat.sc %>%
  group_by(grid_x, grid_y, cell_type) %>%
  summarise(count = n(),
            #            x_center = mean(x),
            #            y_center = mean(y),
```

```
            .groups = 'drop')

total_counts <- dat.sc %>%
  group_by(grid_x, grid_y) %>%
  summarise(total = n(), .groups = 'drop')

proportions <- cell_type_counts %>%
  left_join(total_counts, by = c("grid_x", "grid_y")) %>%
  mutate(proportion = count / total) %>%
  select(grid_x, grid_y, cell_type, proportion)

# Pivot the proportions table for each cell type into columns
proportions_wide <- proportions %>%
  pivot_wider(names_from = cell_type,
              values_from = proportion,
              values_fill = list(proportion = 0))

# Get spot-level data frame
dat.st <- expression_summary <- dat.sc %>%
  group_by(grid_x, grid_y) %>%
  summarise(
    x_center = mean(x),
    y_center = mean(y),
    .groups = 'drop'
  ) %>%
  left_join(proportions_wide, by = c("grid_x", "grid_y")) %>%
  select(x_center, y_center, CT1, CT2, CT3)

# Coordinates matrix
pos.original <- dat.st %>%
  select(x = x_center, y = y_center)
pos.original <- as.matrix(pos.original)
row.names(pos.original) <- paste0("Spot", 1:dim(pos.original)[1])

# Proportion matrix
prop <- dat.st %>%
  select(-c(x_center, y_center))
prop <- as.matrix(prop)
row.names(prop) <- paste0("Spot", 1:dim(prop)[1])

# Expression count matrix
counts <- counts.st
row.names(counts) <- paste0("Gene", 1:dim(counts)[1])
colnames(counts) <- paste0("Spot", 1:dim(counts)[2])
```

#### Rotate the tissue

Define the `rotate_points` function.

```
rotate_points <- function(original_points, angle_degrees) {
  angle_radians <- angle_degrees * (pi / 180)  # Convert degrees to radians
  rotation_matrix <- matrix(c(cos(angle_radians), -sin(angle_radians),
```

```
                              sin(angle_radians),  cos(angle_radians)),
                            nrow = 2, byrow = TRUE)
  rotated_points <- t(rotation_matrix %*% t(original_points))
  output <- as.matrix(rotated_points)
  colnames(output) <- c('x', 'y')
  return(output)
}
```

Set the rotation angle to be 30° or other degrees, rotate the tissue and then scale the new x and y coordinates.

```
# angle degree of rotation
rotate.angle <- 30
# Rotate the original pattern by the given angle degree
pos.rotated.null <- rotate_points(pos.original, angle_degrees = rotate.angle)
# Scale the rotated coordinates
pos.rotated <- pos.rotated.null
pos.rotated[,1] <- (pos.rotated.null[,1] - min(pos.rotated.null)) /
  (max(pos.rotated.null) - min(pos.rotated.null))
pos.rotated[,2] <- (pos.rotated.null[,2] - min(pos.rotated.null)) /
  (max(pos.rotated.null) - min(pos.rotated.null))
```

The original gene expression pattern for Gene 31:

```
gene_expression <- counts.st[31,]
scaled_gene_expression <- (gene_expression - min(gene_expression)) /
  (max(gene_expression) - min(gene_expression))
dat.st.original <- data.frame(x = pos.original[,1],
                              y = pos.original[,2],
                              gene_expression = scaled_gene_expression
                              )
ggplot(dat.st.original, aes(x, y, color = gene_expression)) +
  geom_point() +
  theme_minimal() +
  scale_color_continuous(low = "cornsilk", high = "darkred") +
  theme(legend.title = element_text(size = 11, face = "bold"),
        panel.grid.major = element_blank(),
        panel.grid.minor = element_blank()) +
  labs(color = "Scaled gene expression") +
  guides(color = guide_legend(override.aes = list(size = 4)))
```

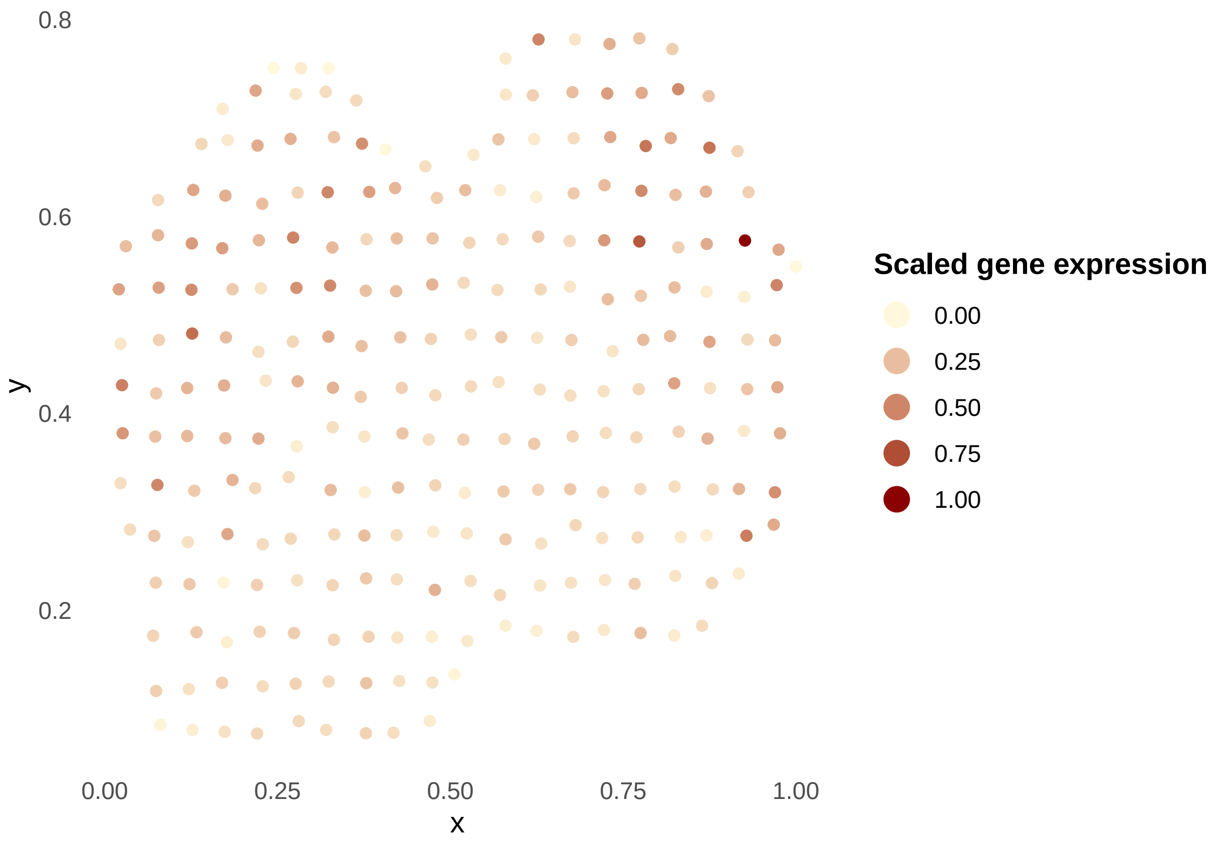


The rotated gene expression pattern for Gene 31:

```
dat.st.rotated <- data.frame(x = pos.rotated[,1],
                              y = pos.rotated[,2],
                              gene_expression = scaled_gene_expression
                              )
ggplot(dat.st.rotated, aes(x, y, color = gene_expression)) +
  geom_point() +
  theme_minimal() +
  scale_color_continuous(low = "cornsilk", high = "darkred") +
  theme(legend.title = element_text(size = 11, face = "bold"),
        panel.grid.major = element_blank(),
        panel.grid.minor = element_blank()) +
  labs(color = "Scaled gene expression") +
  guides(color = guide_legend(override.aes = list(size = 4)))
```

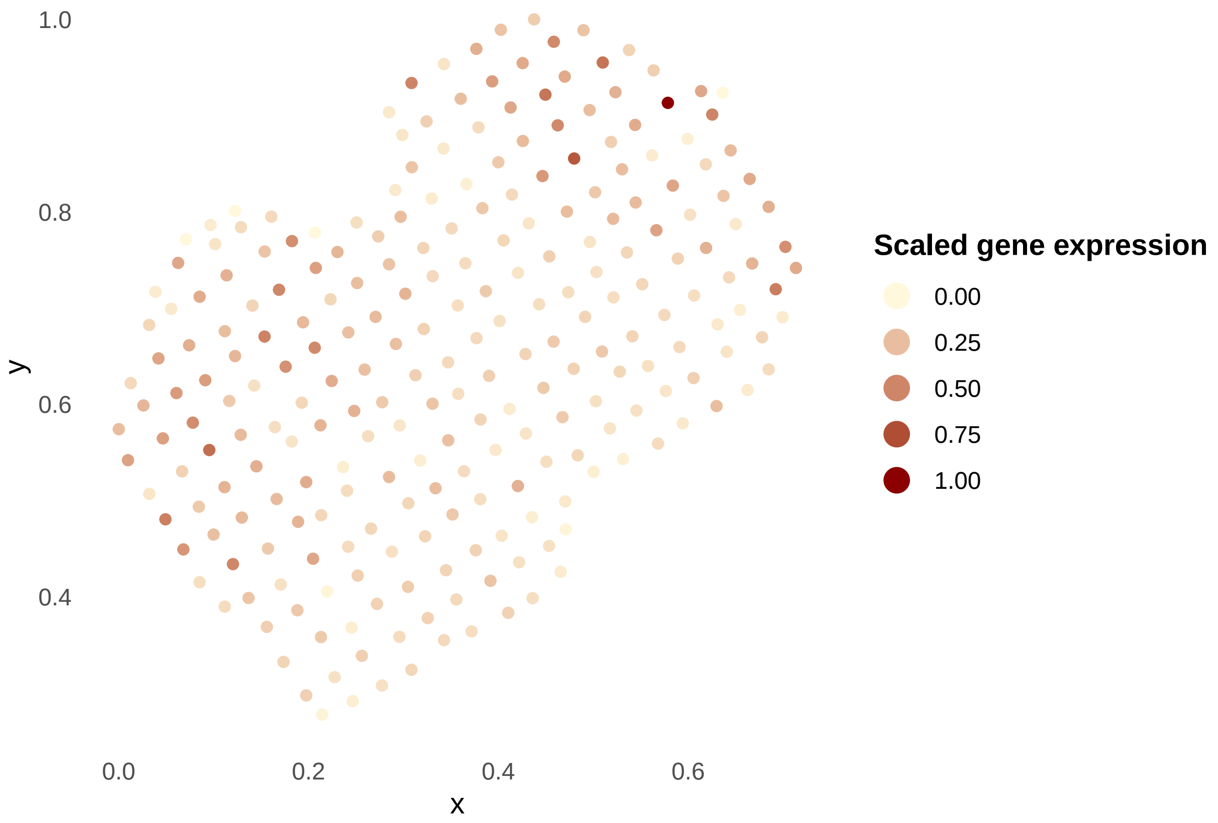


#### CSIDE

Next, we analyzed the data with CSIDE [3] using the data before and after rotation. In theory, the results should be exactly the same given that the relative relationship between gene expressions and their spatial locations remain unchanged. We first created the RCTD [4] objects for both original and rotated patterns.

```
Cell_Types <- as.factor(Cell_Types)
names(Cell_Types) <- paste0("Cell", 1:length(Cell_Types))
# Create the Reference object
RCTD_reference <- Reference(counts = counts.sc,
                            cell_types = Cell_Types, min_UMI = 1)
# Create the SpatialRNA object for the original pattern
RCTD_puck.original <- SpatialRNA(coords = as.data.frame(pos.original),
                                 counts = counts)
# Create the RCTD object for the original pattern
myRCTD.original <- create.RCTD(RCTD_puck.original, RCTD_reference, max_cores = 1)
# Run RCTD for the original pattern
myRCTD.original <- run.RCTD(myRCTD.original, doublet_mode = "full")
# Import the true cell type composition
myRCTD.original <- import_weights(myRCTD.original, weights = prop)

# Create the SpatialRNA object for the rotated pattern
RCTD_puck.rotated <- SpatialRNA(coords = as.data.frame(pos.rotated),
                                counts = counts)
```

```r
# Create the RCTD object for the rotated pattern
myRCTD.rotated <- create.RCTD(RCTD_puck.rotated, RCTD_reference, max_cores = 1)
# Run RCTD for the rotated pattern
myRCTD.rotated <- run.RCTD(myRCTD.rotated, doublet_mode = "full")
# Import the true cell type composition
myRCTD.rotated <- import_weights(myRCTD.rotated, weights = prop)
```

Run CSIDE using `run.CSIDE.nonparam` for both original and rotated patterns.

```r
# Run CSIDE for the original pattern
CSIDE.results.original <- run.CSIDE.nonparam(myRCTD.original, df = 6,
                                             cell_types = paste0('CT',1:3),
                                             gene_threshold = .001,
                                             cell_type_threshold = 10,
                                             fdr = 0.01, doublet_mode = FALSE)
# Run CSIDE for the rotated pattern
CSIDE.results.rotated <- run.CSIDE.nonparam(myRCTD.rotated, df = 6,
                                            cell_types = paste0('CT',1:3),
                                            gene_threshold = .001,
                                            cell_type_threshold = 10,
                                            fdr = 0.01, doublet_mode = FALSE)
```

Then, we checked the significant gene list for cell type 1 (CT1).

```r
print(CSIDE.results.original@de_results$sig_gene_list$CT1)
```

```
##         Z_score   log_fc        se paramindex_best conv        p_val
## Gene36 4.674277 1.852129 0.3962386               3 TRUE 1.474955e-05
## Gene34 3.895407 2.637814 0.6771600               3 TRUE 4.901690e-04
## Gene37 0.000000 0.000000 0.0000000               0 TRUE 6.635030e-04
```

```r
print(CSIDE.results.rotated@de_results$sig_gene_list$CT1)
```

```
##         Z_score    log_fc        se paramindex_best conv        p_val
## Gene36 4.604733 -2.445881 0.5311668               2 TRUE 0.0000206498
## Gene39 3.874766 -2.798021 0.7221137               2 TRUE 0.0005336364
## Gene34 0.000000  0.000000 0.0000000               0 TRUE 0.0005563409
```

We can see that Gene 37 is considered CT1-specific ctSVG before rotation, but no longer significant after rotation, indicating false negative of CSIDE for this gene after rotation. The results keep changing after rotating the coordinates with different angles. This shows that the testing results of CSIDE are not spatial rotation-invariant.

##### spVC

spVC [5] uses the bivariate penalized spline over triangulation (BPST) method to approximate cell type-specific spatial effects, which requires pre-selected boundary points. We preselected the boundary points for this sample pattern and saved as `Sample_stBoundary.rda` (available at https://drive.google.com/drive/folders/1KSxeInbwFswuJdTBxCjMZKc6voz5UWqZ?usp=drive_link). Next we loaded the file and used TriMesh to create trigangulations for the original and rotated patterns.

```r
## For the original pattern
load(file = 'Sample_stBoundary.rds') # Load the pre-selected boundary points
# TriMesh is used to create triangulation
Tr.cell.original <- TriMesh(mouse.cerebellum.boundary.original, n = 2)
V.original <- as.matrix(Tr.cell.original$V)
Tr.original <- as.matrix(Tr.cell.original$Tr)
# The boundary points of rotated pattern
mouse.cerebellum.boundary.rotated <- rotate_points(mouse.cerebellum.boundary.original,
                                                   angle_degrees = rotate.angle)
#points(mouse.cerebellum.boundary.rotated, type = "l", col = "blue")
Tr.cell.rotated <- TriMesh(mouse.cerebellum.boundary.rotated, n = 2)
V.rotated.null <- as.matrix(Tr.cell.rotated$V)
# Scale the V coordinate matrix
V.rotated <- V.rotated.null
V.rotated[,1] <- (V.rotated.null[,1] - min(pos.rotated.null)) /
  (max(pos.rotated.null) - min(pos.rotated.null))
V.rotated[,2] <- (V.rotated.null[,2] - min(pos.rotated.null)) /
  (max(pos.rotated.null) - min(pos.rotated.null))
Tr.rotated <- as.matrix(Tr.cell.rotated$Tr)
```

Then, spVC was run for both data before and after rotation.

```r
# Fit the spVC model for the original pattern
spVC.results.original <- test.spVC(Y = counts, X = prop, S = pos.original,
                                   V = V.original, Tr = Tr.original,
                                   para.cores = 1, filter.min.nonzero = 5)
```

```
## spVC model will use  94.42231 % of the original data.
## Conducting tests for 50  genes.
## Model 2: Conducting tests for 0 genes.
```

```r
# Fit the spVC model for the rotated pattern
spVC.results.rotated <- test.spVC(Y = counts, X = prop, S = pos.rotated,
                                  V = V.rotated, Tr = Tr.rotated,
                                  para.cores = 1, filter.min.nonzero = 5)
```

```
## spVC model will use  94.42231 % of the original data.
## Conducting tests for 50  genes.
## Model 2: Conducting tests for 1 genes.
```

We checked the testing results for one of the CT1-specific ctSVG. Here, we selected Gene 31 as an example.

```r
# spVC
print(spVC.results.original$results.constant$Gene31$p.value)
```

```
##       beta_0      beta_X1      beta_X2      beta_X3      gamma_0
## 7.515357e-45 5.653109e-01          NaN 1.086993e-01 1.075567e-11
```

```r
print(spVC.results.rotated$results.constant$Gene31$p.value)
```

```
##       beta_0      beta_X1      beta_X2      beta_X3      gamma_0
## 5.796108e-39 1.346266e-02 8.961199e-02          NaN 1.431855e-12
```

Gene 31 did not pass the stage 1 test before rotation, since none of p-values for testing the cell-type-associated spatially constant effects `beta_X1, beta_X2, beta_X3` was significant at the 0.05 level, even though the residual spatial effect `gamma_0` was significant (p-value=1.08E-11). On the other hand, it passed the stage 1 test after rotation with significant residual spatial effect `gamma_0` and CT1-associated spatially constant effect `beta_X1`. This shows that the testing results of spVC are not invariant to spatial rotation.

##### CTSV

Finally, we performed CTSV [6] on the data before and after rotation.

```r
spe.original <- SpatialExperiment(assay = counts[,which(colSums(counts) != 0)],
                                  colData = pos.original,
                                  spatialCoordsNames = c('x', 'y'))
spe.rotated <- SpatialExperiment(assay = counts[,which(colSums(counts) != 0)],
                                 colData = pos.rotated,
                                 spatialCoordsNames = c('x', 'y'))

CTSV.results.original <- CTSV(spe.original, W = prop, num_core = 1)
CTSV.results.rotated <- CTSV(spe.rotated, W = prop, num_core = 1)
```

We checked the significant gene list for cell type 1 (CT1).

```r
print(svGene(CTSV.results.original$qval, 0.05)$SVGene[[1]])
```

```
## [1] "Gene13" "Gene32" "Gene34" "Gene40"
```

```r
print(svGene(CTSV.results.rotated$qval, 0.05)$SVGene[[1]])
```

```
## [1] "Gene32" "Gene34" "Gene37" "Gene38" "Gene39" "Gene40"
```

We can see that Gene 37, Gene 38 and Gene 39 were considered as CT1-specific ctSVGs in the rotated pattern, but not in the original pattern, indicating that CTSV is not invariant to spatial rotation.

##### Summary

In summary, we simulated 10 CT1-specific ctSVGs and applied the three methods to the data before and after a 30-degree spatial rotation. We observed inconsistencies in the testing results before and after rotation, even though the results should have remained unchanged. Furthermore, as the rotation angle varied, different testing outcomes were observed. These inconsistent results highlight that these methods are unreliable for ctSVG detection and should be avoided in ctSVG analysis.
