## Supplementary File II for "STANCE: a unified statistical model to detect cell-type-specific spatially variable genes in spatial transcriptomics"

#### 1 Details of STANCE

##### 1.1 The model

Our goal is to identify genes that exhibit spatial expression patterns, referred to as spatially variable genes (SVGs), specific to certain cell types. Suppose we have spatial transcriptomics expression data for $m$ genes from $n$ spots (or pixels) of a 2D tissue, with their spatial locations denoted as $\mathbf{s} = (s_{i1}, s_{i2})_{n\times 2}$, $i = 1, \cdots, n$. The original gene expression count data of $n$ spots are collected and normalized through various methods to yield continuous gene expression data, denoted as $\mathbf{y} = (y_1, \cdots, y_n)^T$. Furthermore, we assume that all cells in this tissue belong to $K$ cell types, and for each spot, the cell type compositions are estimated using existing cell type deconvolution methods, either reference-based such as RCTD [1] or [2] or reference-free such as STdeconvolve [3], denoted by $\mathbf{\Pi} = (\pi_{i1}, \cdots, \pi_{iK})_{n\times K}$, $i = 1, \cdots, n$.

With this information, we establish a variance component model[4] to elucidate the relationship between gene expressions and spatial locations, i.e.,

$$\mathbf{y}(\mathbf{s}) = \mathbf{X}(\mathbf{s})\boldsymbol{\beta} + \boldsymbol{\gamma}(\mathbf{s}) + \boldsymbol{\varepsilon}(\mathbf{s}), \tag{1}$$

where $\mathbf{X}(\mathbf{s})$ is a $n \times p$-dimensional design matrix for covariates and $\boldsymbol{\beta} = (\beta_1, \cdots, \beta_p)^T$ being a $p$-dimensional vector of associated coefficients (in the default case, $\mathbf{X}(\mathbf{s})\boldsymbol{\beta}$ contains only the intercept); $\boldsymbol{\gamma}(\mathbf{s})$ is a random spatial effect component, and $\boldsymbol{\varepsilon}(\mathbf{s}) = (\varepsilon_1, \cdots, \varepsilon_n)^T$ is a $n$-dimensional vector of random effects of the error term, following a multivariate normal distribution $MVN(\mathbf{0}, \sigma_\varepsilon^2\mathbf{I}_n)$. Model(1) is the SVG detection model commonly assumed in the literature, e.g., in SpatialDE, SPARK or nnSVG (ref). In order to model the cell-type-specific effect, we further decompose the random effect term $\boldsymbol{\gamma}$ into $K$ components, i.e.,

$$\boldsymbol{\gamma}(\mathbf{s}) = \boldsymbol{\pi}_1 \odot \boldsymbol{\gamma}_1(\mathbf{s}) + \cdots + \boldsymbol{\pi}_K \odot \boldsymbol{\gamma}_K(\mathbf{s}), \tag{2}$$

where $\boldsymbol{\gamma}_k(\mathbf{s})$ is an $n$-dimensional vector of spatial random effects contributed by cell type $k$, $k = 1, \cdots, K$; $\boldsymbol{\pi}_k = (\pi_{1k}, \cdots, \pi_{nk})^T$ is the $k$-th column of $\mathbf{\Pi}$, whose $(i, k)$-th element $\pi_{ik}$ is the proportion of a specific cell type $k$ in spot $i$, $i = 1, \cdots, n$; $\odot$ is the Hamadard element-wise product. We assume that cell-type spatial random effects are independent of each other and each cell-type spatial random effect follows a multivariate normal distribution,

$$\boldsymbol{\gamma}_k(\mathbf{s}) = (\gamma_k(\mathbf{s}_1), \cdots, \gamma_k(\mathbf{s}_n))^T \sim MVN(\mathbf{0}, \tau_k \mathbf{K}), k = 1, \cdots, K, \tag{3}$$

where $\mathbf{K}$ is an $n \times n$-dimensional kernel matrix capturing the spatial similarity between spots; $\tau_k$ is the variance component of spatial effect corresponding to cell type $k$. Combining model(1) and (2), we have

$$\mathbf{y}(\mathbf{s}) = \mathbf{X}(\mathbf{s})\boldsymbol{\beta} + \boldsymbol{\pi}_1 \odot \boldsymbol{\gamma}_1(\mathbf{s}) + \cdots + \boldsymbol{\pi}_K \odot \boldsymbol{\gamma}_K(\mathbf{s}) + \boldsymbol{\varepsilon}(\mathbf{s}). \tag{4}$$

which is the final STANCE model for cell-type-specific SVG detection. The covariance of $\mathbf{y}$ is given by

$$\mathbf{V} = \mathrm{Cov}(\mathbf{y}) = \sum_{k=1}^{K} \tau_k \boldsymbol{\Pi}_k \mathbf{K} \boldsymbol{\Pi}_k^T + \sigma_\varepsilon^2 \mathbf{I}_n = \sum_{k=1}^{K} \tau_k \boldsymbol{\Sigma}_k + \sigma_\varepsilon^2 \mathbf{I}_n, \tag{5}$$

where $\boldsymbol{\Pi}_k = \mathrm{diag}\{\boldsymbol{\pi}_k\}$ and $\boldsymbol{\Sigma}_k = \boldsymbol{\Pi}_k \mathbf{K} \boldsymbol{\Pi}_k^T$ for $k = 1, \cdots, K$.

#### 1.2 The estimation

Denote $\boldsymbol{\tau} = (\tau_1, \cdots, \tau_K)^T$, $\boldsymbol{\phi} = (\boldsymbol{\tau}^T, \sigma_\varepsilon^2)^T$, and $\boldsymbol{\theta} = (\boldsymbol{\beta}^T, \boldsymbol{\phi}^T)^T$, then the likelihood function for $\boldsymbol{\theta}$ is

$$L(\boldsymbol{\theta}; \mathbf{y}) = (2\pi)^{-\frac{n}{2}} |\mathbf{V}|^{-\frac{1}{2}} \exp\{-\frac{1}{2}(\mathbf{y} - \mathbf{X}\boldsymbol{\beta})^T \mathbf{V}^{-1} (\mathbf{y} - \mathbf{X}\boldsymbol{\beta})\}. \tag{6}$$

Then, the log-likelihood function for $\boldsymbol{\theta}$ is given by

$$l(\boldsymbol{\theta}; \mathbf{y}) = -\frac{n}{2} \log(2\pi) - \frac{1}{2} \log|\mathbf{V}| - \frac{1}{2}(\mathbf{y} - \mathbf{X}\boldsymbol{\beta})^T \mathbf{V}^{-1} (\mathbf{y} - \mathbf{X}\boldsymbol{\beta}) \tag{7}$$

Take the first derivative of the log likelihood function with respect to $\boldsymbol{\beta}$ and set it to 0, we obtain the restricted maximum likelihood (REML) estimator as[5, 6]

$$\hat{\boldsymbol{\beta}} = (\mathbf{X}^T \mathbf{V}^{-1} \mathbf{X})^{-1} \mathbf{X}^T \mathbf{V}^{-1} \mathbf{y} \tag{8}$$

Profiling out $\boldsymbol{\beta}$ and taking logarithm gives us the restricted log likelihood function for $\boldsymbol{\phi}$,

$$\begin{aligned} l^R(\boldsymbol{\phi}; \mathbf{y}) &\propto -\frac{1}{2} \log |\mathbf{V}| - \frac{1}{2} \log |\mathbf{X}^T \mathbf{V}^{-1} \mathbf{X}| - \frac{1}{2}(\mathbf{y} - \mathbf{X}\hat{\boldsymbol{\beta}})^T \mathbf{V}^{-1} (\mathbf{y} - \mathbf{X}\hat{\boldsymbol{\beta}}) \\ &\propto -\frac{1}{2} \log |\mathbf{V}| - \frac{1}{2} \log |\mathbf{X}^T \mathbf{V}^{-1} \mathbf{X}| - \frac{1}{2} \mathbf{y}^T \mathbf{P} \mathbf{y}, \end{aligned} \tag{9}$$

where $\mathbf{P} = \mathbf{V}^{-1} - \mathbf{V}^{-1}\mathbf{X}(\mathbf{X}^T \mathbf{V}^{-1} \mathbf{X})^{-1} \mathbf{X}^T \mathbf{V}^{-1}$.

Then, the score vector of $\boldsymbol{\phi}$ is given by

$$\mathbf{S}(\boldsymbol{\phi}; \mathbf{y}) = \left(S(\tau_1; \mathbf{y}), \cdots, S(\tau_K; \mathbf{y}), S(\sigma_\varepsilon^2; \mathbf{y})\right)^T, \tag{10}$$

where

$$
\begin{aligned}
S(\tau_k;\mathbf{y}) &= \frac{\partial l^R(\boldsymbol{\phi};\mathbf{y})}{\partial \tau_k} \\
&= -\frac{1}{2}\mathrm{tr}(\mathbf{V}^{-1}\boldsymbol{\Sigma}_k) + \frac{1}{2}\mathrm{tr}\big[(\mathbf{X}^T\mathbf{V}^{-1}\mathbf{X})^{-1}\mathbf{X}^T\mathbf{V}^{-1}\boldsymbol{\Sigma}_k\mathbf{V}^{-1}\mathbf{X}\big] + \frac{1}{2}\mathbf{y}'\mathbf{P}\boldsymbol{\Sigma}_k\mathbf{P}\mathbf{y} \\
&= -\frac{1}{2}\mathrm{tr}\Big\{\big[\mathbf{V}^{-1} - \mathbf{V}^{-1}\mathbf{X}(\mathbf{X}^T\mathbf{V}^{-1}\mathbf{X})^{-1}\mathbf{X}^T\mathbf{V}^{-1}\big]\boldsymbol{\Sigma}_k\Big\} + \frac{1}{2}\mathbf{y}'\mathbf{P}\boldsymbol{\Sigma}_k\mathbf{P}\mathbf{y} \\
&= -\frac{1}{2}\mathrm{tr}(\mathbf{P}\boldsymbol{\Sigma}_k) + \frac{1}{2}\mathbf{y}'\mathbf{P}\boldsymbol{\Sigma}_k\mathbf{P}\mathbf{y}, \qquad (11)\\
S(\sigma_\varepsilon^2;\mathbf{y}) &= \frac{\partial l^R(\boldsymbol{\phi};\mathbf{y})}{\partial \sigma_\varepsilon^2} \\
&= -\frac{1}{2}\mathrm{tr}(\mathbf{V}^{-1}\mathbf{I}_n) + \frac{1}{2}\mathrm{tr}\big[(\mathbf{X}^T\mathbf{V}^{-1}\mathbf{X})^{-1}\mathbf{X}^T\mathbf{V}^{-1}\mathbf{I}_n\mathbf{V}^{-1}\mathbf{X}\big] + \frac{1}{2}\mathbf{y}'\mathbf{P}\mathbf{I}_n\mathbf{P}\mathbf{y} \\
&= -\frac{1}{2}\mathrm{tr}\Big\{\big[\mathbf{V}^{-1} - \mathbf{V}^{-1}\mathbf{X}(\mathbf{X}^T\mathbf{V}^{-1}\mathbf{X})^{-1}\mathbf{X}^T\mathbf{V}^{-1}\big]\mathbf{I}_n\Big\} + \frac{1}{2}\mathbf{y}'\mathbf{P}\boldsymbol{\Sigma}_k\mathbf{P}\mathbf{y} \\
&= -\frac{1}{2}\mathrm{tr}(\mathbf{P}\mathbf{I}_n) + \frac{1}{2}\mathbf{y}'\mathbf{P}\mathbf{I}_n\mathbf{P}\mathbf{y}. \qquad (12)
\end{aligned}
$$

We further compute the second derivatives of the REML log-likelihood

$$
\begin{aligned}
\frac{\partial^2 l^R(\boldsymbol{\phi};\mathbf{y})}{\partial\tau_k\partial\tau_l} &= \frac{1}{2}\mathrm{tr}(\mathbf{P}\boldsymbol{\Sigma}_l\mathbf{P}\boldsymbol{\Sigma}_k) - \frac{1}{2}\mathbf{y}'\mathbf{P}\boldsymbol{\Sigma}_l\mathbf{P}\boldsymbol{\Sigma}_k\mathbf{P}\mathbf{y} - \frac{1}{2}\mathbf{y}'\mathbf{P}\boldsymbol{\Sigma}_k\mathbf{P}\boldsymbol{\Sigma}_l\mathbf{P}\mathbf{y} \\
&= \frac{1}{2}\mathrm{tr}(\mathbf{P}\boldsymbol{\Sigma}_l\mathbf{P}\boldsymbol{\Sigma}_k) - \mathbf{y}'\mathbf{P}\boldsymbol{\Sigma}_k\mathbf{P}\boldsymbol{\Sigma}_l\mathbf{P}\mathbf{y} \\
\frac{\partial^2 l^R(\boldsymbol{\phi};\mathbf{y})}{\partial\tau_k\partial\sigma_\varepsilon^2} &= \frac{1}{2}\mathrm{tr}(\mathbf{P}\mathbf{I}_n\mathbf{P}\boldsymbol{\Sigma}_k) - \frac{1}{2}\mathbf{y}'\mathbf{P}\mathbf{I}_n\mathbf{P}\boldsymbol{\Sigma}_k\mathbf{P}\mathbf{y} - \frac{1}{2}\mathbf{y}'\mathbf{P}\boldsymbol{\Sigma}_k\mathbf{P}\mathbf{I}_n\mathbf{P}\mathbf{y} \\
&= \frac{1}{2}\mathrm{tr}(\mathbf{P}\mathbf{I}_n\mathbf{P}\boldsymbol{\Sigma}_k) - \mathbf{y}'\mathbf{P}\boldsymbol{\Sigma}_k\mathbf{P}\mathbf{I}_n\mathbf{P}\mathbf{y} \\
\frac{\partial^2 l^R(\boldsymbol{\phi};\mathbf{y})}{\partial\sigma_\varepsilon^2\partial\tau_k} &= \frac{1}{2}\mathrm{tr}(\mathbf{P}\boldsymbol{\Sigma}_k\mathbf{P}\mathbf{I}_n) - \frac{1}{2}\mathbf{y}'\mathbf{P}\boldsymbol{\Sigma}_k\mathbf{P}\mathbf{I}_n\mathbf{P}\mathbf{y} - \frac{1}{2}\mathbf{y}'\mathbf{P}\mathbf{I}_n\mathbf{P}\boldsymbol{\Sigma}_k\mathbf{P}\mathbf{y} \\
&= \frac{1}{2}\mathrm{tr}(\mathbf{P}\boldsymbol{\Sigma}_k\mathbf{P}\mathbf{I}_n) - \mathbf{y}'\mathbf{P}\boldsymbol{\Sigma}_k\mathbf{P}\mathbf{I}_n\mathbf{P}\mathbf{y} \\
\frac{\partial^2 l^R(\boldsymbol{\phi};\mathbf{y})}{\partial^2\sigma_\varepsilon^2} &= \frac{1}{2}\mathrm{tr}(\mathbf{P}\mathbf{I}_n\mathbf{P}\mathbf{I}_n) - \frac{1}{2}\mathbf{y}'\mathbf{P}\mathbf{I}_n\mathbf{P}\mathbf{I}_n\mathbf{P}\mathbf{y} - \frac{1}{2}\mathbf{y}'\mathbf{P}\mathbf{I}_n\mathbf{P}\mathbf{I}_n\mathbf{P}\mathbf{y} \\
&= \frac{1}{2}\mathrm{tr}(\mathbf{P}\mathbf{I}_n\mathbf{P}\mathbf{I}_n) - \mathbf{y}'\mathbf{P}\mathbf{I}_n\mathbf{P}\mathbf{I}_n\mathbf{P}\mathbf{y}
\end{aligned}
$$

Since we have

$$
\begin{aligned}
\mathbb{E}(\mathbf{y}'\mathbf{P}\boldsymbol{\Sigma}_k\mathbf{P}\boldsymbol{\Sigma}_l\mathbf{P}\mathbf{y}) &= \mathrm{tr}(\mathbf{P}\boldsymbol{\Sigma}_l\mathbf{P}\boldsymbol{\Sigma}_k) \\
\mathbb{E}(\mathbf{y}'\mathbf{P}\boldsymbol{\Sigma}_k\mathbf{P}\mathbf{I}_n\mathbf{P}\mathbf{y}) &= \mathrm{tr}(\mathbf{P}\mathbf{I}_n\mathbf{P}\boldsymbol{\Sigma}_k) \\
\mathbb{E}(\mathbf{y}'\mathbf{P}\mathbf{I}_n\mathbf{P}\mathbf{I}_n\mathbf{P}\mathbf{y}) &= \mathrm{tr}(\mathbf{P}\mathbf{I}_n\mathbf{P}\mathbf{I}_n)
\end{aligned}
$$

then the average information matrix[7] is given by

$$
\begin{aligned}
\mathbf{AI}(\boldsymbol{\theta};\mathbf{y}) &= \begin{bmatrix}
\frac{1}{2}\Big[\frac{\partial^2 l^R(\boldsymbol{\phi};\mathbf{y})}{\partial\tau_1^2} + \mathbb{E}\big(\frac{\partial^2 l^R(\boldsymbol{\phi};\mathbf{y})}{\partial\tau_1^2}\big)\Big] & \cdots & \frac{1}{2}\Big[\frac{\partial^2 l^R(\boldsymbol{\phi};\mathbf{y})}{\partial\tau_1\partial\tau_K} + \mathbb{E}\big(\frac{\partial^2 l^R(\boldsymbol{\phi};\mathbf{y})}{\partial\tau_1\partial\tau_K}\big)\Big] & \frac{1}{2}\Big[\frac{\partial^2 l^R(\boldsymbol{\phi};\mathbf{y})}{\partial\tau_1\partial\sigma_\varepsilon^2} + \mathbb{E}\big(\frac{\partial^2 l^R(\boldsymbol{\phi};\mathbf{y})}{\partial\tau_1\partial\sigma_\varepsilon^2}\big)\Big] \\
\vdots & \ddots & \vdots & \vdots \\
\frac{1}{2}\Big[\frac{\partial^2 l^R(\boldsymbol{\phi};\mathbf{y})}{\partial\tau_K\partial\tau_1} + \mathbb{E}\big(\frac{\partial^2 l^R(\boldsymbol{\phi};\mathbf{y})}{\partial\tau_K\partial\tau_1}\big)\Big] & \cdots & \frac{1}{2}\Big[\frac{\partial^2 l^R(\boldsymbol{\phi};\mathbf{y})}{\partial\tau_K^2} + \mathbb{E}\big(\frac{\partial^2 l^R(\boldsymbol{\phi};\mathbf{y})}{\partial\tau_K^2}\big)\Big] & \frac{1}{2}\Big[\frac{\partial^2 l^R(\boldsymbol{\phi};\mathbf{y})}{\partial\tau_K\partial\sigma_\varepsilon^2} + \mathbb{E}\big(\frac{\partial^2 l^R(\boldsymbol{\phi};\mathbf{y})}{\partial\tau_K\partial\sigma_\varepsilon^2}\big)\Big] \\
\frac{1}{2}\Big[\frac{\partial^2 l^R(\boldsymbol{\phi};\mathbf{y})}{\partial\sigma_\varepsilon^2\partial\tau_1} + \mathbb{E}\big(\frac{\partial^2 l^R(\boldsymbol{\phi};\mathbf{y})}{\partial\sigma_\varepsilon^2\partial\tau_1}\big)\Big] & \cdots & \frac{1}{2}\Big[\frac{\partial^2 l^R(\boldsymbol{\phi};\mathbf{y})}{\partial\sigma_\varepsilon^2\partial\tau_K} + \mathbb{E}\big(\frac{\partial^2 l^R(\boldsymbol{\phi};\mathbf{y})}{\partial\sigma_\varepsilon^2\partial\tau_K}\big)\Big] & \frac{1}{2}\Big[\frac{\partial^2 l^R(\boldsymbol{\phi};\mathbf{y})}{\partial} + \mathbb{E}\big(\frac{\partial^2 l^R(\boldsymbol{\phi};\mathbf{y})}{\partial\sigma_\varepsilon^4}\big)\Big]
\end{bmatrix} \\
&= \frac{1}{2}\begin{bmatrix}
\mathbf{y}'\mathbf{P}\boldsymbol{\Sigma}_1\mathbf{P}\boldsymbol{\Sigma}_1\mathbf{P}\mathbf{y} & \cdots & \mathbf{y}'\mathbf{P}\boldsymbol{\Sigma}_1\mathbf{P}\boldsymbol{\Sigma}_K\mathbf{P}\mathbf{y} & \mathbf{y}'\mathbf{P}\boldsymbol{\Sigma}_1\mathbf{P}\mathbf{I}_n\mathbf{P}\mathbf{y} \\
\vdots & \ddots & \vdots & \vdots \\
\mathbf{y}'\mathbf{P}\boldsymbol{\Sigma}_K\mathbf{P}\boldsymbol{\Sigma}_1\mathbf{P}\mathbf{y} & \cdots & \mathbf{y}'\mathbf{P}\boldsymbol{\Sigma}_K\mathbf{P}\boldsymbol{\Sigma}_K\mathbf{P}\mathbf{y} & \mathbf{y}'\mathbf{P}\boldsymbol{\Sigma}_K\mathbf{P}\mathbf{I}_n\mathbf{P}\mathbf{y} \\
\mathbf{y}'\mathbf{P}\mathbf{I}_n\mathbf{P}\boldsymbol{\Sigma}_1\mathbf{P}\mathbf{y} & \cdots & \mathbf{y}'\mathbf{P}\mathbf{I}_n\mathbf{P}\boldsymbol{\Sigma}_K\mathbf{P}\mathbf{y} & \mathbf{y}'\mathbf{P}\mathbf{I}_n\mathbf{P}\mathbf{I}_n\mathbf{P}\mathbf{y}
\end{bmatrix}.
\end{aligned} \tag{13}
$$

We can apply the Newton's method for the estimates of variance components $\boldsymbol{\phi}$ by the following iterative update[7]

$$\boldsymbol{\phi}^{(t+1)} = \boldsymbol{\phi}^{(t)} + \mathbf{AI}(\boldsymbol{\theta};\mathbf{y})\mathbf{S}(\boldsymbol{\phi}^{(t)};\mathbf{y}), \tag{14}$$

and further obtain the estimates of $\boldsymbol{\beta}$. In practice, we implement the "lmm_aireml" function in the R package 'gaston'[8] to refine the estimates more efficiently.

#### 1.3 Hypothesis testing

##### 1.3.1 Overall score test for detecting utSVGs

To detect utSVGs, we consider the following null and alternative hypotheses

$$\begin{cases} H_0^{(1)} : \tau_1 = \cdots = \tau_K = 0 \\ H_1^{(1)} : \text{at least one parameter is not zero} \end{cases} \tag{15}$$

Under the null hypothesis $H_0^1$, the null model is given by

$$\mathbf{y}(\mathbf{s}) = \mathbf{X}(\mathbf{s})\boldsymbol{\beta} + \boldsymbol{\varepsilon}(\mathbf{s}) \sim MVN(\mathbf{X}\boldsymbol{\beta}, \sigma_\varepsilon^2\mathbf{I}_n). \tag{16}$$

Then, by fitting the above null model, we obtain $\hat{\boldsymbol{\beta}} = (\mathbf{X}^T\mathbf{X})^{-1}\mathbf{X}^T\mathbf{y}$ and $\hat{\sigma}_\varepsilon^2 = \frac{(\mathbf{y}-\mathbf{X}\hat{\boldsymbol{\beta}})^T(\mathbf{y}-\mathbf{X}\hat{\boldsymbol{\beta}})}{n}$. Since $\mathbf{V}_0 = \sigma_\varepsilon^2\mathbf{I}_n$, then the corresponding $\mathbf{P}_0$ matrix is given by

$$\mathbf{P}_0 = \mathbf{V}_0^{-1} - \mathbf{V}_0^{-1}\mathbf{X}(\mathbf{X}^T\mathbf{V}_0^{-1}\mathbf{X})^{-1}\mathbf{X}^T\mathbf{V}_0^{-1} = \frac{1}{\sigma_\varepsilon^2}\big[\mathbf{I}_n - \mathbf{X}(\mathbf{X}^T\mathbf{X})^{-1}\mathbf{X}^T\big]. \tag{17}$$

According to formula(11), the REML score function for $\tau_k$ under the null is

$$S_0(\tau_k) = -\frac{1}{2}\text{tr}(\mathbf{P}_0\boldsymbol{\Sigma}_k) + \frac{1}{2}\mathbf{y}^T\mathbf{P}_0\boldsymbol{\Sigma}_k\mathbf{P}_0\mathbf{y} \tag{18}$$

The REML version of score test statistic for $\boldsymbol{\tau} = (\tau_1, \cdots, \tau_K)^T$ under $H_0^1$ is

$$U^{(1)}(\boldsymbol{\tau}) = \sum_{k=1}^{K}\frac{1}{2}\mathbf{y}'\mathbf{P}_0\boldsymbol{\Sigma}_k\mathbf{P}_0\mathbf{y} = \frac{1}{2}\mathbf{y}^T\mathbf{P}_0\boldsymbol{\Sigma}\mathbf{P}_0\mathbf{y}, \tag{19}$$

where $\mathbf{\Sigma} = \sum_{k=1}^{K} \mathbf{\Sigma}_k$, which is a quadratic form of $\mathbf{y}$ following a mixture of chi-square distributions under the null[9, 10]. Specifically, plugging in $\mathbf{P}_0$ in (17) to the score statistic in (19), we get

$$U^{(1)}(\boldsymbol{\tau}) = \frac{1}{2}\mathbf{y}^T\mathbf{P}_0\mathbf{\Sigma}\mathbf{P}_0\mathbf{y} = \frac{1}{2\sigma_\varepsilon^4}\mathbf{y}^T\tilde{\mathbf{P}}_0\mathbf{\Sigma}\tilde{\mathbf{P}}_0\mathbf{y}, \tag{20}$$

where $\tilde{\mathbf{P}}_0 = \mathbf{I}_n - \mathbf{X}(\mathbf{X}^T\mathbf{X})^{-1}\mathbf{X}^T$ is an idempotent projection matrix.

To compute the corresponding p-value, the distribution of the test statistic $U^{(1)}(\boldsymbol{\tau})$ can be approximated by a scaled chi-square distribution $a\chi_g^2$ through the Satterthwaite approximation method[11]. Specifically, the mean $e$ and variance $v$ of $U^{(1)}(\boldsymbol{\tau})$ are given by

$$\begin{cases} e &= \mathbb{E}\left[U^{(1)}(\boldsymbol{\tau})\right] = \frac{1}{2\sigma_\varepsilon^2}\text{tr}(\tilde{\mathbf{P}}_0\mathbf{\Sigma}) \\ v &= \text{Var}\left[U^{(1)}(\boldsymbol{\tau})\right] = \frac{1}{2\sigma_\varepsilon^4}\text{tr}\left[(\tilde{\mathbf{P}}_0\mathbf{\Sigma})(\tilde{\mathbf{P}}_0\mathbf{\Sigma})\right] \end{cases} \tag{21}$$

Then, we match them with the expected moments under the scaled chi-square distribution $a\chi_g^2$ to calculate the scale parameter $a$ and the degree of freedom $g$, i.e.,

$$\begin{cases} e &= \mathbb{E}\left[U^{(1)}(\boldsymbol{\tau})\right] \equiv \mathbb{E}(a\chi_g^2) = ag \\ v &= \text{Var}\left[U^{(1)}(\boldsymbol{\tau})\right] \equiv \text{Var}(a\chi_g^2) = 2a^2g \end{cases} \tag{22}$$

Solve the above equations(22), we get

$$\begin{cases} a &= \frac{v}{2e} \\ g &= \frac{2e^2}{v} \end{cases} \tag{23}$$

Given that the true value of $\sigma_\varepsilon^2$ is unknown, replacing $\sigma_\varepsilon^2$ in (21) by its estimate $\hat{\sigma}_\varepsilon^2$ gives the estimates $\hat{e}$ and $\hat{v}$, and further the estimates $\hat{a}$ and $\hat{v}$ by

$$\begin{cases} \hat{a} &= \frac{\hat{v}}{2\hat{e}} \\ \hat{g} &= \frac{2\hat{e}^2}{\hat{v}} \end{cases} \tag{24}$$

To account for the fact that $\boldsymbol{\beta}$ and $\sigma_\varepsilon^2$ are estimated by their REML estimates under the null model, a bias correction is introduced for estimating $a$ and $g$[9]. Suppose that $\boldsymbol{\beta}$ is known, the Fisher information matrix for $\boldsymbol{\phi} = (\tau_1^2, \cdots, \tau_K^2, \sigma_\varepsilon^2)^T$ can be partitioned as

$$\mathcal{I}(\tau_1, \cdots, \tau_K, \sigma_\varepsilon^2) = \begin{bmatrix} \mathcal{I}_{\boldsymbol{\tau},\boldsymbol{\tau}} & \mathcal{I}_{\boldsymbol{\tau},\sigma} \\ \mathcal{I}_{\boldsymbol{\tau},\sigma}^T & \mathcal{I}_{\sigma,\sigma} \end{bmatrix}, \tag{25}$$

where $\mathcal{I}_{\boldsymbol{\tau},\boldsymbol{\tau}}$ is the expected Fisher information for $\boldsymbol{\tau}$, whose $(i,j)$-th entry is $\frac{1}{2\sigma_\varepsilon^4}\text{tr}\left[(\tilde{\mathbf{P}}_0\mathbf{\Sigma}_i)(\tilde{\mathbf{P}}_0\mathbf{\Sigma}_j)\right]$, $i,j = 1, \cdots, K$; $\mathcal{I}_{\sigma,\sigma} = \frac{1}{2\sigma_\varepsilon^4}\text{tr}(\tilde{\mathbf{P}}_0\mathbf{I}_n\tilde{\mathbf{P}}_0\mathbf{I}_n) = \frac{1}{2\sigma_\varepsilon^4}\text{tr}(\tilde{\mathbf{P}}_0)$ is the expected Fisher information for $\sigma_\varepsilon^2$; and

$$\mathcal{I}_{\boldsymbol{\tau},\sigma} = \frac{1}{2\sigma_\varepsilon^4}\left[\text{tr}(\tilde{\mathbf{P}}_0\mathbf{I}_n\tilde{\mathbf{P}}_0\mathbf{\Sigma}_1), \cdots, \text{tr}(\tilde{\mathbf{P}}_0\mathbf{I}_n\tilde{\mathbf{P}}_0\mathbf{\Sigma}_K)\right]^T = \frac{1}{2\sigma_\varepsilon^4}\left[\text{tr}(\tilde{\mathbf{P}}_0\mathbf{\Sigma}_1), \cdots, \text{tr}(\tilde{\mathbf{P}}_0\mathbf{\Sigma}_K)\right]^T. \tag{26}$$

Replacing $\sigma_\varepsilon^2$ by its REML estimator $\hat{\sigma}_\varepsilon^2$ for every partition of $\mathcal{I}(\tau_1, \cdots, \tau_K, \sigma_\varepsilon^2)$ leads to $I_{\boldsymbol{\tau},\boldsymbol{\tau}}$, $I_{\boldsymbol{\tau},\sigma}$ and $I_{\sigma,\sigma}$.

Notice that $v = \frac{1}{2\sigma_\varepsilon^4}\mathrm{tr}\big[(\tilde{\mathbf{P}}_0\boldsymbol{\Sigma})(\tilde{\mathbf{P}}_0\boldsymbol{\Sigma})\big] = \mathbf{1}_n^T\mathcal{I}_{\tau,\tau}\mathbf{1}_n$, then we estimate $v$ by replacing $\mathcal{I}_{\tau,\tau}$ with the efficient information

$$\tilde{v} = \mathbf{1}_n^T\tilde{I}_{\boldsymbol{\tau},\boldsymbol{\tau}}\mathbf{1}_n, \tag{27}$$

where $\tilde{I}_{\boldsymbol{\tau},\boldsymbol{\tau}} = I_{\boldsymbol{\tau},\boldsymbol{\tau}} - I_{\boldsymbol{\tau},\sigma}I_{\sigma,\sigma}^{-1}I_{\boldsymbol{\tau},\sigma}^T$ is the efficient information for $\boldsymbol{\tau}$.

Then, the bias-corrected estimates for $a$ and $g$ are given by

$$\begin{cases} \tilde{a} &= \frac{\tilde{v}}{2\hat{e}} \\ \tilde{g} &= \frac{2\hat{e}^2}{\tilde{v}} \end{cases}. \tag{28}$$

The corresponding p-value at significance level $\alpha$ can be computed by $\mathbb{P}(\frac{U^{(1)}(\boldsymbol{\tau})}{\tilde{a}} > \chi^2_{\tilde{g},\alpha})$.

##### 1.3.2 Cell-type-specific individual score test to detect ctSVGs

For a specific cell type $l$, to detect its associated ctSVGs, we consider the following hypotheses,

$$\begin{cases} H_0^{(2)}: \tau_l = 0 \\ H_1^{(2)}: \tau_l > 0 \end{cases} \tag{29}$$

Under the null hypothesis $H_0^{(2)}$, we obtain the reduced model

$$\mathbf{y}(\mathbf{s}) = \mathbf{X}(\mathbf{s})\boldsymbol{\beta} + \sum_{k\neq l}^{n}\boldsymbol{\Pi}_k\boldsymbol{\gamma}_k(\mathbf{s}) + \boldsymbol{\varepsilon}(\mathbf{s}), \tag{30}$$

The variance of $\mathbf{y}$ under this null is given by $\mathbf{V}_{-l} = \sum_{k\neq l}^K \tau_k\boldsymbol{\Pi}_k\mathbf{K}\boldsymbol{\Pi}_k + \sigma_\varepsilon^2\mathbf{I}_n$, and the associated $\mathbf{P}$ matrix is

$$\mathbf{P}_{-l} = \mathbf{V}_{-l}^{-1} - \mathbf{V}_{-l}^{-1}\mathbf{X}(\mathbf{X}^T\mathbf{V}_{-l}^{-1}\mathbf{X})^{-1}\mathbf{X}^T\mathbf{V}_{-l}^{-1}.$$

Therefore, the REML version of the score test statistic for $\tau_l$ under $H_0^{(2)}$ is

$$U^{(2)}(\tau_l) = \frac{1}{2}\mathbf{y}^T\mathbf{P}_{-l}\boldsymbol{\Sigma}_l\mathbf{P}_{-l}\mathbf{y}, \tag{31}$$

which is also a quadratic form of $\mathbf{y}$ following a mixture of chi-square distributions under the null[9]. Similarly, we use the Satterthwaite method to approximate the distribution of $U^{(2)}(\tau_l)$ by a scaled chi-square distribution $a_l\chi^2_{g_l}$.

In this case, the mean and the variance of $U^{(2)}(\tau_l)$ are

$$\begin{cases} e_1 &= \mathbb{E}\big[U^{(2)}(\tau_l)\big] = \frac{1}{2}\mathrm{tr}(\mathbf{P}_{-l}\boldsymbol{\Sigma}_l) \\ v_1 &= \mathrm{Var}\big[U^{(2)}(\tau_l)\big] = \frac{1}{2}\mathrm{tr}\big[(\mathbf{P}_{-l}\boldsymbol{\Sigma}_l)(\mathbf{P}_{-l}\boldsymbol{\Sigma}_l)\big] \end{cases} \tag{32}$$

Given that $\sigma_\varepsilon^2$ and $\boldsymbol{\tau}_{-l} = (\tau_1, \cdots, \tau_{l-1}, \tau_{l+1}, \cdots, \tau_K)^T$ are unknown, we replace them in $\mathbf{P}_{-l}$ by their REML estimates under the reduced model(30) to obtain $\hat{\mathbf{P}}_{-l}$ and further the estimates $\hat{e}_l$ and $\hat{v}_l$. To account for this fact, we will also perform bias correction for the test statistic $U^{(2)}(\tau_l)$[9].

We realign the elements in $\boldsymbol{\phi}_l$ to get $\boldsymbol{\phi}_l = (\sigma_\varepsilon^2, \tau_1, \cdots, \tau_{l-1}, \tau_{l+1}, \cdots, \tau_K, \tau_l)^T$. Suppose $\boldsymbol{\beta}$ is known, the Fisher information matrix for $\boldsymbol{\phi}_l$ can be partitioned as

$$\mathcal{I}(\boldsymbol{\phi}_l) = \begin{bmatrix} \mathcal{I}_{-l,-l} & \mathcal{I}_{-l,l} \\ \mathcal{I}_{-l,l}^T & \mathcal{I}_{l,l} \end{bmatrix}. \tag{33}$$

where

$$\mathcal{I}_{-l,-l} = \frac{1}{2}\begin{bmatrix} \mathrm{tr}(\mathbf{P}_{-l}\mathbf{P}_{-l}) & \mathrm{tr}(\mathbf{P}_{-l}\mathbf{P}_{-l}\mathbf{\Sigma}_1) & \cdots & \mathrm{tr}(\mathbf{P}_{-l}\mathbf{P}_{-l}\mathbf{\Sigma}_K) \\ \mathrm{tr}(\mathbf{P}_{-l}\mathbf{\Sigma}_1\mathbf{P}_{-l}) & \mathrm{tr}(\mathbf{P}_{-l}\mathbf{\Sigma}_1\mathbf{P}_{-l}\mathbf{\Sigma}_1) & \cdots & \mathrm{tr}(\mathbf{P}_{-l}\mathbf{\Sigma}_1\mathbf{P}_{-l}\mathbf{\Sigma}_K) \\ \vdots & \vdots & \ddots & \vdots \\ \mathrm{tr}(\mathbf{P}_{-l}\mathbf{\Sigma}_K\mathbf{P}_{-l}) & \mathrm{tr}(\mathbf{P}_{-l}\mathbf{\Sigma}_K\mathbf{P}_{-l}\mathbf{\Sigma}_1) & \cdots & \mathrm{tr}(\mathbf{P}_{-l}\mathbf{\Sigma}_K\mathbf{P}_{-l}\mathbf{\Sigma}_K), \end{bmatrix} \tag{34}$$

is the expected Fisher information for $\boldsymbol{\tau}_{-l}$;

$$\mathcal{I}_{l,l} = \frac{1}{2}\mathrm{tr}(\mathbf{P}_{-l}\mathbf{\Sigma}_l\mathbf{P}_{-l}\mathbf{\Sigma}_l) \tag{35}$$

is the expected Fisher information for $\sigma_\varepsilon^2$; and

$$\mathcal{I}_{-l,l} = \frac{1}{2}\left[\mathrm{tr}(\mathbf{P}_{-l}\mathbf{\Sigma}_l\mathbf{P}_{-l}), \mathrm{tr}(\mathbf{P}_{-l}\mathbf{\Sigma}_l\mathbf{P}_{-l}\mathbf{\Sigma}_1), \cdots, \mathrm{tr}(\mathbf{P}_{-l}\mathbf{\Sigma}_l\mathbf{P}_{-l}\mathbf{\Sigma}_K)\right]^T. \tag{36}$$

Replacing $\mathbf{P}_{-l}$ by $\hat{\mathbf{P}}_{-l}$ evaluated at $\hat{\sigma}_\varepsilon^2$, $\hat{\boldsymbol{\tau}}_{-l}$ leads to $I(\boldsymbol{\phi}_l)$ under the null model. Notice that $v_l = \mathcal{I}_{l,l}$, then we estimate $v_l$ by the efficient information

$$\tilde{I}_{l,l} = I_{l,l} - I_{-l,l}^T I_{-l,-l}^{-1} I_{-l,l} \equiv \tilde{v}_l. \tag{37}$$

Then, the bias-corrected estimates for $a_l$ and $g_l$ are given by

$$\begin{cases} \tilde{a}_l & = \tilde{v}_l / 2\hat{e}_l \\ \tilde{g}_l & = 2\hat{e}_l^2 / \tilde{v}_l \end{cases}. \tag{38}$$

The corresponding p-value at significance level $\alpha$ can be computed by $\mathbb{P}(\frac{U^{(2)}(\tau_l)}{\tilde{a}_l} > \chi^2_{\tilde{g}_l,\alpha})$.

#### 1.4 Kernel matrix

The kernel matrix, which represents the spatial correlation pattern of spots within the target tissue, is crucial for parameter estimation and statistical inference. In our approach, we employ a Gaussian kernel defined as $\mathbf{K}(\mathbf{s}_i, \mathbf{s}_j) = \exp\{-\frac{|\mathbf{s}_i - \mathbf{s}_j|^2}{h}\}$, where $|\mathbf{s}_i - \mathbf{s}_j|$ denotes the Euclidean distance between spots $\mathbf{s}_i$ and $\mathbf{s}_j$, and $h$ is the bandwidth. The Euclidean distance is spatial rotation invariant, and the STANCE model links the gene expression pattern and spatial location through the Gaussian kernel. Thus, not like CTSV, C-SIDE, and spVG, the estimation and testing results are invariant to spatial rotation and translation.

To determine the appropriate bandwidth $h$, we adhere to the method proposed by spatialPCA[12]. Specifically, for datasets with 5000 spots or fewer, non-parametric Sheather-Jones' bandwidths[13] are computed for each gene, and the median of these values is used as the common bandwidth $h$; for datasets with more than 5000 spots, Silverman's rule of thumb[14] is modified to calculate a bandwidth for each gene, and the median of these gene-specific bandwidths is used as the common bandwidth $h$. One can also apply the cosine kernel with different parameters and get different p-values, then these p-values can be combined with the Cauchy combination rule as implemented in SPARK.

### 2 Additional details on real data analysis

#### 2.1 Quality control and cell type deconvolution

The human HER2+ breast cancer tumor dataset[15] can be found at https://zenodo.org, and we used sample H1 as an example, which contains gene expression count data for 15,030 genes over 613 spots. Following the paper, we applied Stereoscope[16], a reference-based deconvolution method, to get the cell type composition of each spot. We used the major tier annotation with 8 cell types, including myeloid cells, T cells, B cells, epithelial cells, plasma cells, endothelial cells, cancer-associated fibroblasts (CAFs), and Perivascular like cells (PVL cells). We then removed 21 genes identified as ring-pattern technical artifacts in the original study following the work of Stereoscope[16] and low-expressed genes that do not express in more than 10% spots, resulting in 10,053 genes.

The human kidney cancer dataset can be found at https://data.mendeley.com. We focused on the tumor core sample of patient PD47171, which contains expression count data for 36,601 genes across 3,008 spots, we used CARD[2], a reference-based cell type deconvolution tool, to obtain the cell type compositions. Specifically, we followed the default quality control procedure of CARD to remove low-expressed genes and spots. We obtained the cell type proportions for 12 cell types, including B cells, plasma cells, T cells, natural killer (NK) cells, endothelial cells (EC), renal cell carcinoma (RCC) cells, non-proximal tubule epithelial (Epi_non-PT) cells, proximal tubule epithelial (Epi_PT) cells, fibrobalast cells, myeloid cells, plasmacytoid dendritic cells (pDC), as well as mast cells, across 2,917 spots. We then removed mitochondrial genes and low-expressed genes that do not express at more than 10% spots, resulting in 7,270 genes.

The mouse olfactory bulb (MOB) dataset [17] can be found at https://www.spatialresearch.org. We focused on MOB replicate #8, which consists of expression count data for 15,928 genes across 262 pixels (spots). We then deconvolved the MOB data using STdeconvolve[3], a reference-free and unsupervised cell type deconvolution tool. Before cell type deconvolution, we followed the procedure of the original work of STdeconvolve to clean the dataset, resulting in 7,365 genes and 260 spots.

#### 2.2 Gene set enrichment analyses

The gene set enrichment analyses were conducted using the "gProfiler2" package for the human breast cancer and kidney cancer datasets[18]. We used the 'gost' function to assess the significance of pathways associated with each group of ctSVGs, based on GO, KEGG, and Reactome gene sets. Specifically, we adapted the default "g_SCS" algorithm for multiple testing correction and set significance level to be 0.05.

#### Supplementary figures

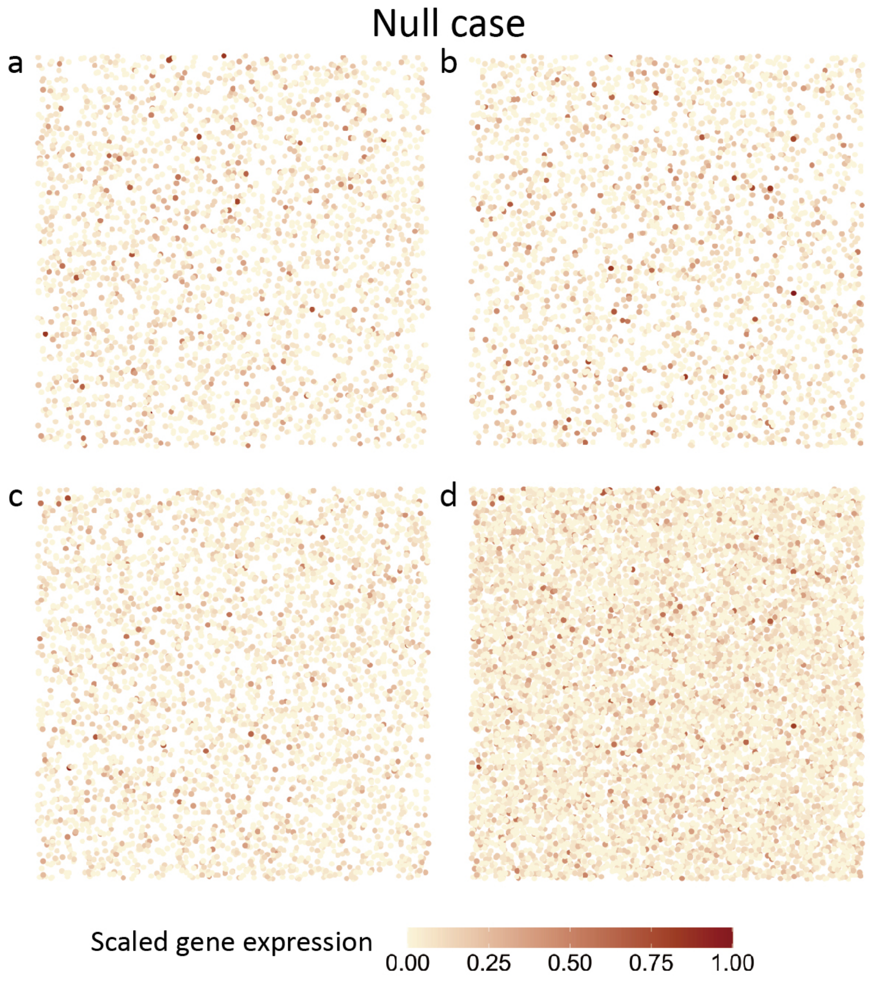


Figure S1: **The spatial expression patterns of a representative non-spatial gene (both non-SVG and non-ctSVG).**

**a.** The spatial expression pattern of cell type 1 shows a random spatial pattern and hence is not a ctSVG.

**b.** The spatial expression pattern of cell type 2 shows a random spatial pattern and hence is not a ctSVG.

**c.** The spatial expression pattern of cell type 3 shows a random spatial pattern and hence is not a ctSVG.

**d.** The spatial expression pattern of the three combined cell types shows a random spatial pattern and hence is not an SVG.

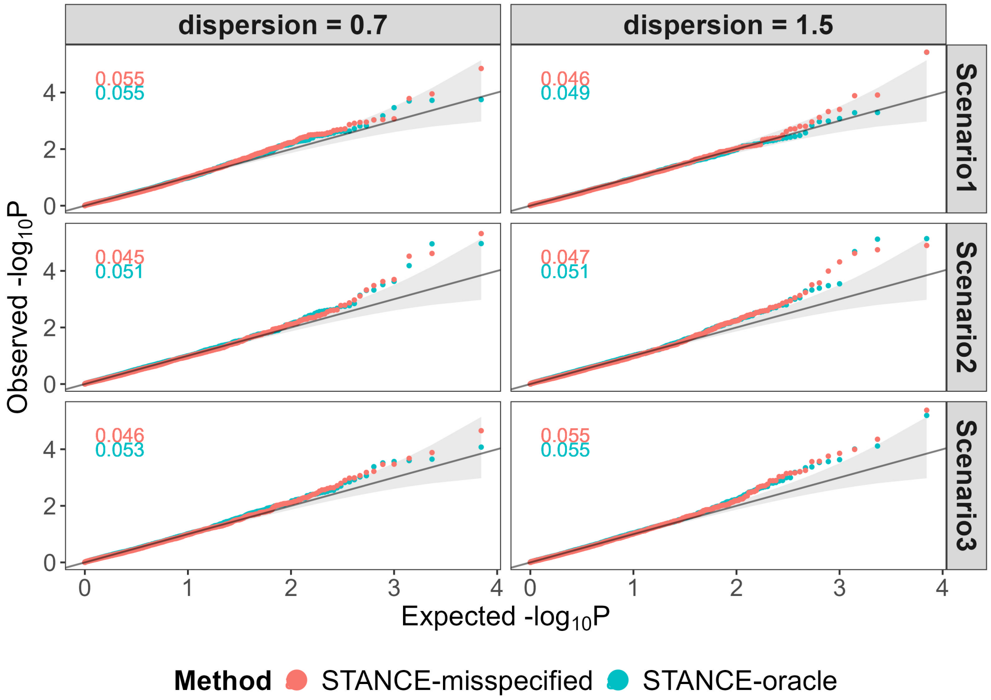


Figure S2: **Simulation results to assess the type I error control for the overall test in Simulation 1 with mis-specified cell type compositions.** The Q-Q plots of the observed $-\log_{10} P$ against the expected $-\log_{10} P$ for different methods are displayed across various dispersion parameters and scenarios. Each sub-figure also includes the empirical type I error rate of different methods, shown in the top left corner, with colors matching those in the legend. When the cell type compositions are miss-deconvolved, the STANCE overall test still controls the type I error well.

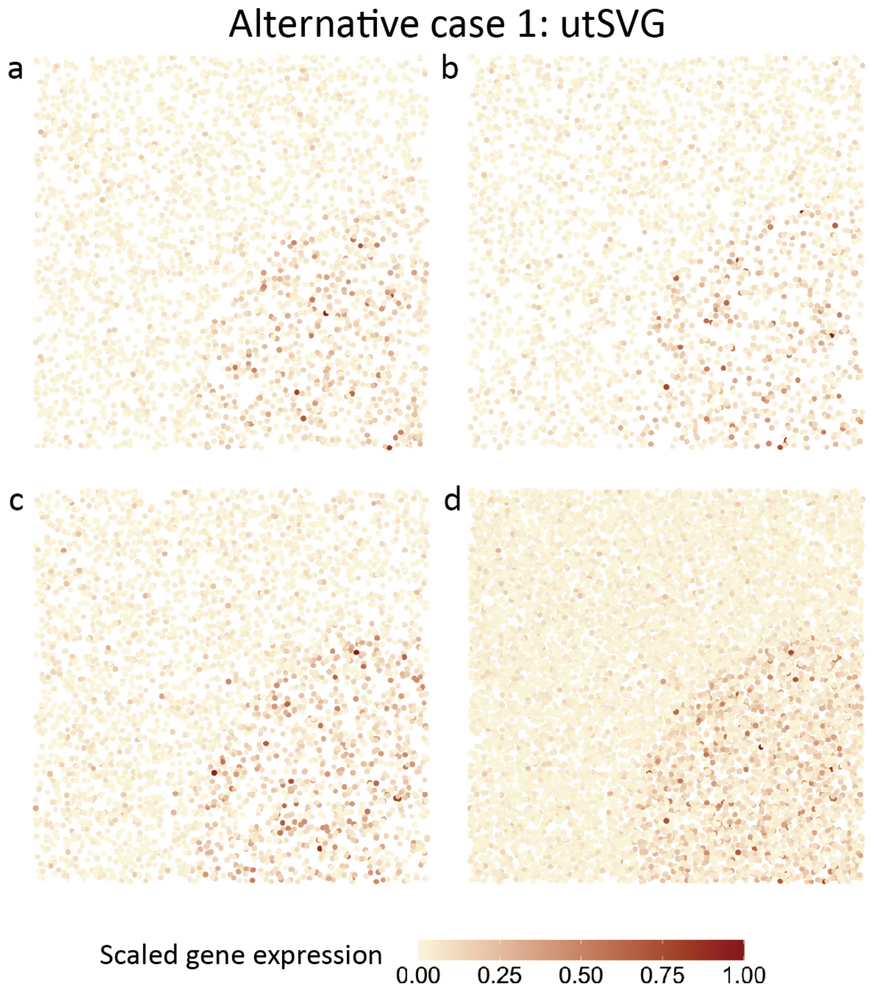


Figure S3: **The spatial patterns of a representative utSVG's expression.** **a-d** show the expression pattern for cell type 1-3 and the combined expression pattern, respectively. The gene is a ctSVG and SVG, hence a utSVG.

**a.** Cell type 1-specific pattern displaying scaled gene expression for cells in cell type 1. The gene expression in domain $D2$ is higher than that in the other domains. Hence, it is a cell type 1 ctSVG.

**b.** Cell type 2-specific pattern displaying scaled gene expression for cells in cell type 2. The gene expression in domain $D2$ is higher than that in the other domains. Hence, it is a cell type 2 ctSVG.

**c.** Cell type 3-specific pattern displaying scaled gene expression for cells in cell type 3. The gene expression in domain $D2$ is higher than that in the other domains. Hence, it is a cell type 3 ctSVG.

**d.** Combined pattern displaying scaled gene expression combining all three cell types. The gene expression in domain $D2$ is higher than that in the other domains. Hence, it is an SVG.

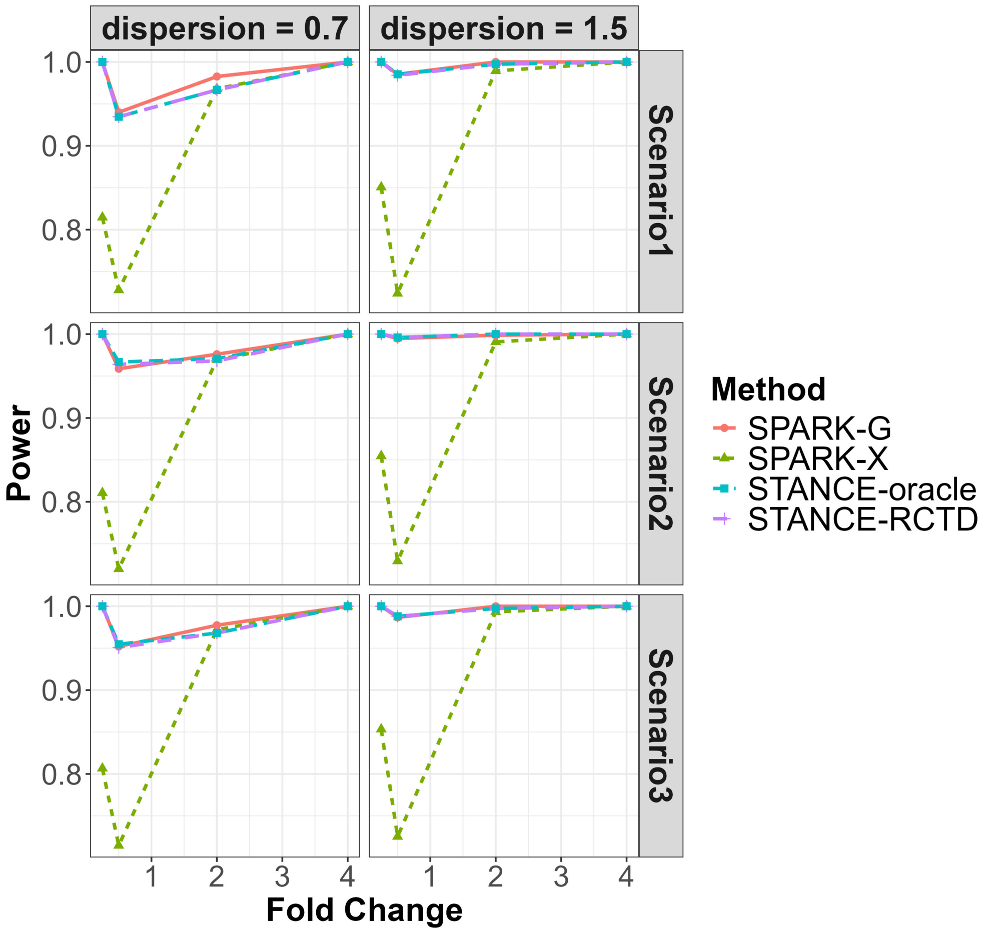


Figure S4: **Simulation results to assess the power of the overall test under Alternative Case 1 in Simulation 1.** Simulation 1 Alternative Case 1 involves both SVGs and ctSVGs. The plots display power values against the fold changes in gene expression for different methods, across various dispersion parameters and cell type composition scenarios, under the significance level of 0.05. In this simulation, STANCE-RCTD and STANCE-oracle perform similarly to SPARK-G and better than SPARK-X.

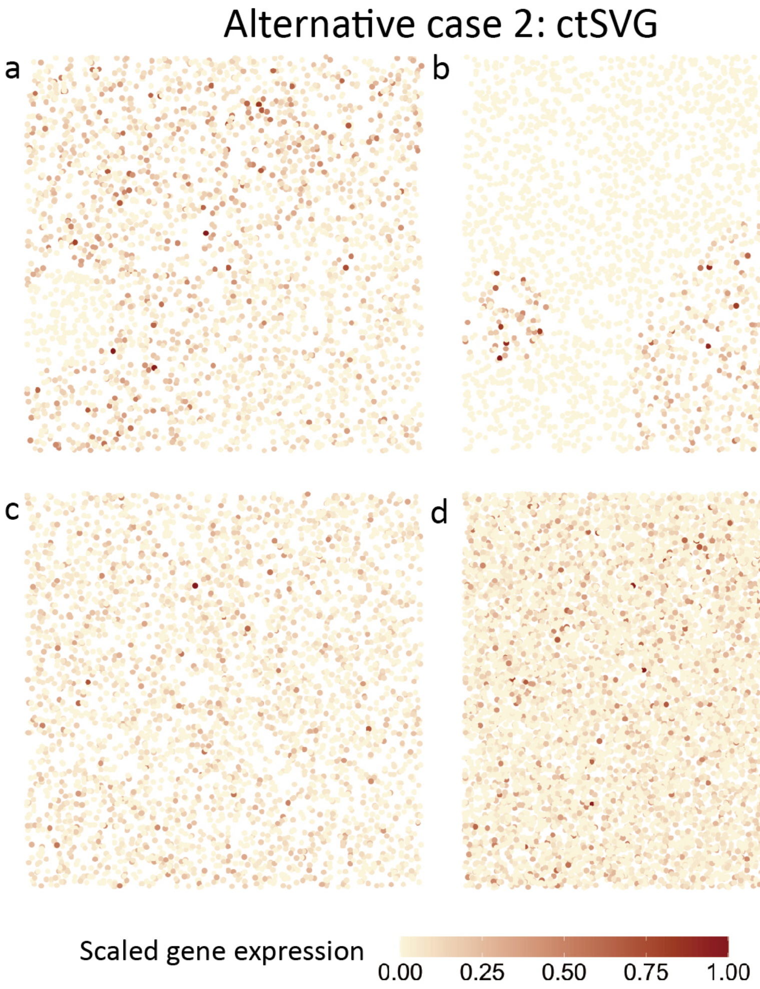


Figure S5: **The spatial expression patterns of a representative ctSVG.** The gene is a cell type 1 and 2 ctSVG but not an SVG and cell type 3 ctSVG.

**a.** Cell type 1-specific pattern displaying scaled gene expression for cells in cell type 1. The gene does not express in domain $D3$. Hence, it is a cell type 1-specific ctSVG.

**b.** Cell type 2-specific pattern displaying scaled gene expression for cells in cell type 2. The gene does not express in domain $D1$. Hence, it is a cell type 2-specific ctSVG.

**c.** Cell type 3-specific pattern displaying scaled gene expression for cells in cell type 3. This representative gene displays random spatial pattern from the perspective of cell type 3. Hence, it is not a cell type 3-specific ctSVG.

**d.** Combined pattern displaying scaled gene expression for all cells. This representative gene displays random spatial pattern from the combined perspective. Hence, it is not an SVG.

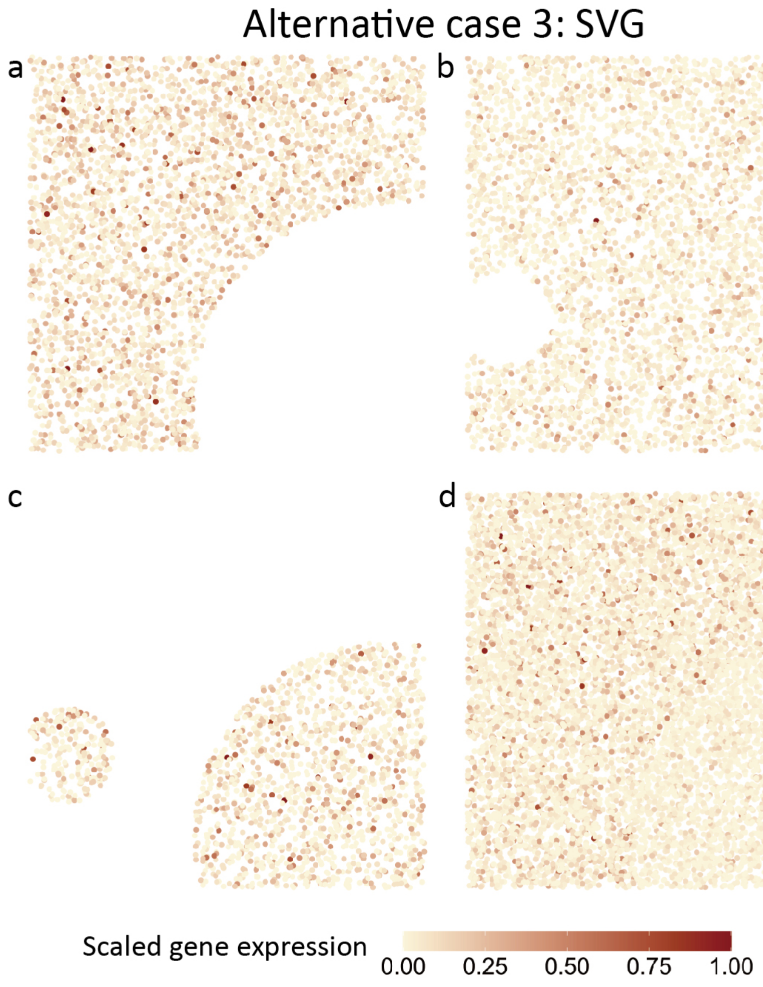


Figure S6: **The spatial expression patterns of a representative SVG.**

**a.** Cell type 1-specific pattern displaying scaled gene expression for cells in cell type 1. This gene displays random spatial pattern in domain D1 and D3, but no expression in D2. Hence, it is not a cell type 1-specific ctSVG.

**b.** Cell type 2-specific pattern displaying scaled gene expression for cells in cell type 2. This gene displays random spatial pattern in domain D1 and D2, but no expression in D3. Hence, it is not a cell type 2-specific ctSVG.

**c.** Cell type 3-specific pattern displaying scaled gene expression for cells in cell type 3. This gene displays random spatial pattern in domain D2 and D3 but not in D1. Hence, it is not a cell type 3-specific ctSVG.

**d.** Combined pattern displaying scaled gene expression in all cells. The gene expression in domain $D2$ is lower than that in the other domains. Hence, it is an SVG.

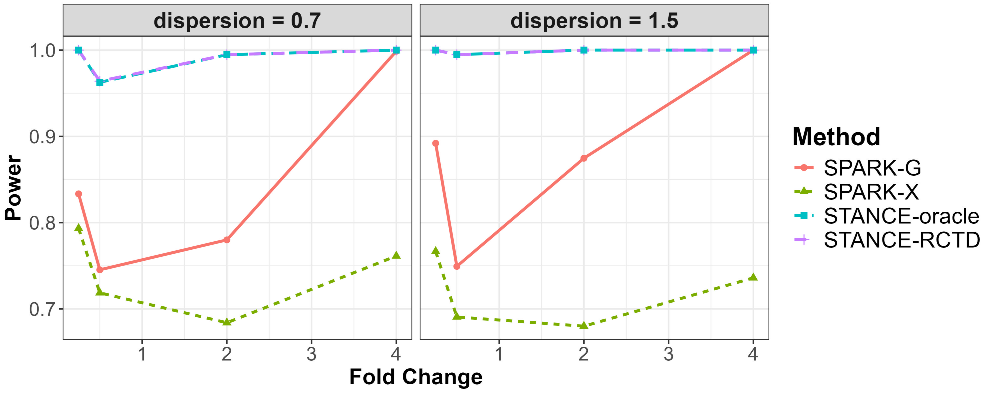


Figure S7: **Simulation results to assess the power of the overall test under Alternative Case 3 in Simulation 1.** Simulation 1 alternative Case 3 involves only SVGs. The plots display power values against the fold changes in gene expression for different methods, across various dispersion parameters, under the significance level of 0.05.

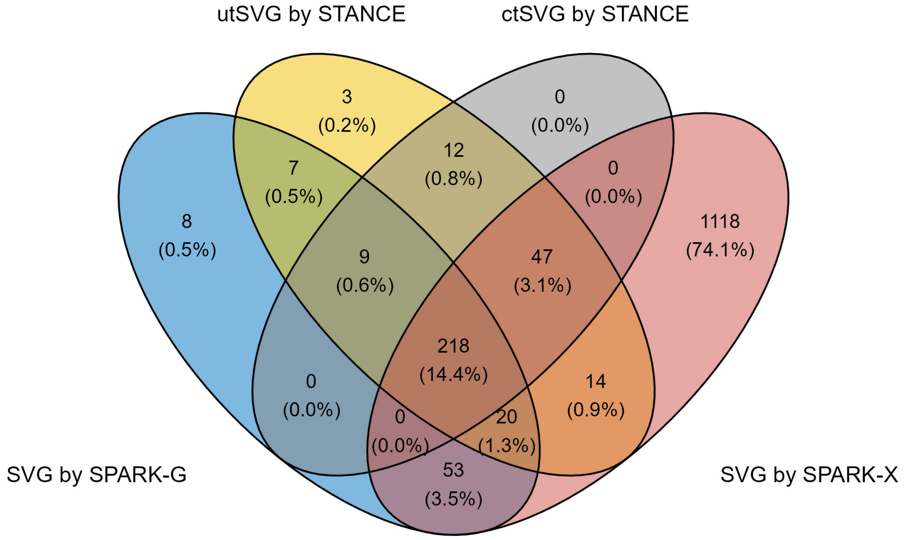


Figure S8: **The Venn diagram for genes identified by different methods in the human breast cancer dataset.** The Venn diagram shows the logical relationship between sets of genes identified by STANCE, SPARK-G, and SPARK-X. The STANCE overall test identified 330 utSVGs (a mixture of SVGs and ctSVGs), with p-values adjusted by the Benjamini-Yekutieli method with an FDR of 0.05. SPARK-G identified 315 SVGs, of which 254 were also identified by the STANCE overall test. SPARK-X identified 1,470 SVGs, with 299 overlapping with those detected by STANCE. For the 330 utSVGs detected by the STANCE overall test, 286 ctSVGs were identified across all 8 cell types by the STANCE cell-type-specific test.

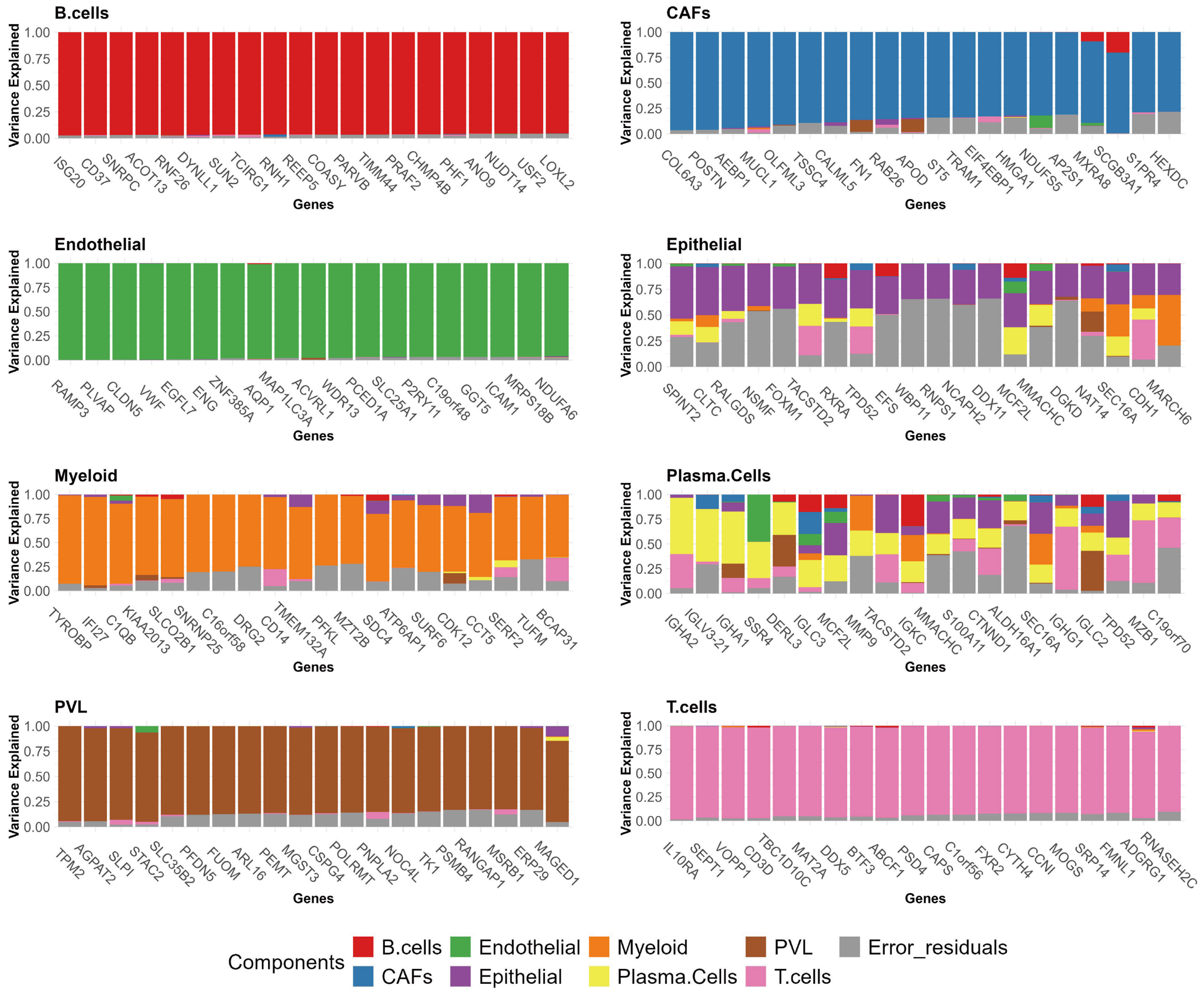


Figure S9: **The stacked variance plots for each cell type in the human breast cancer dataset.** Displayed are the top 20 significant ctSVGs in each cell type. For each gene, the stacked bar plots display the proportion of variance explained by the 8 cell type-specific spatial effects and random error.

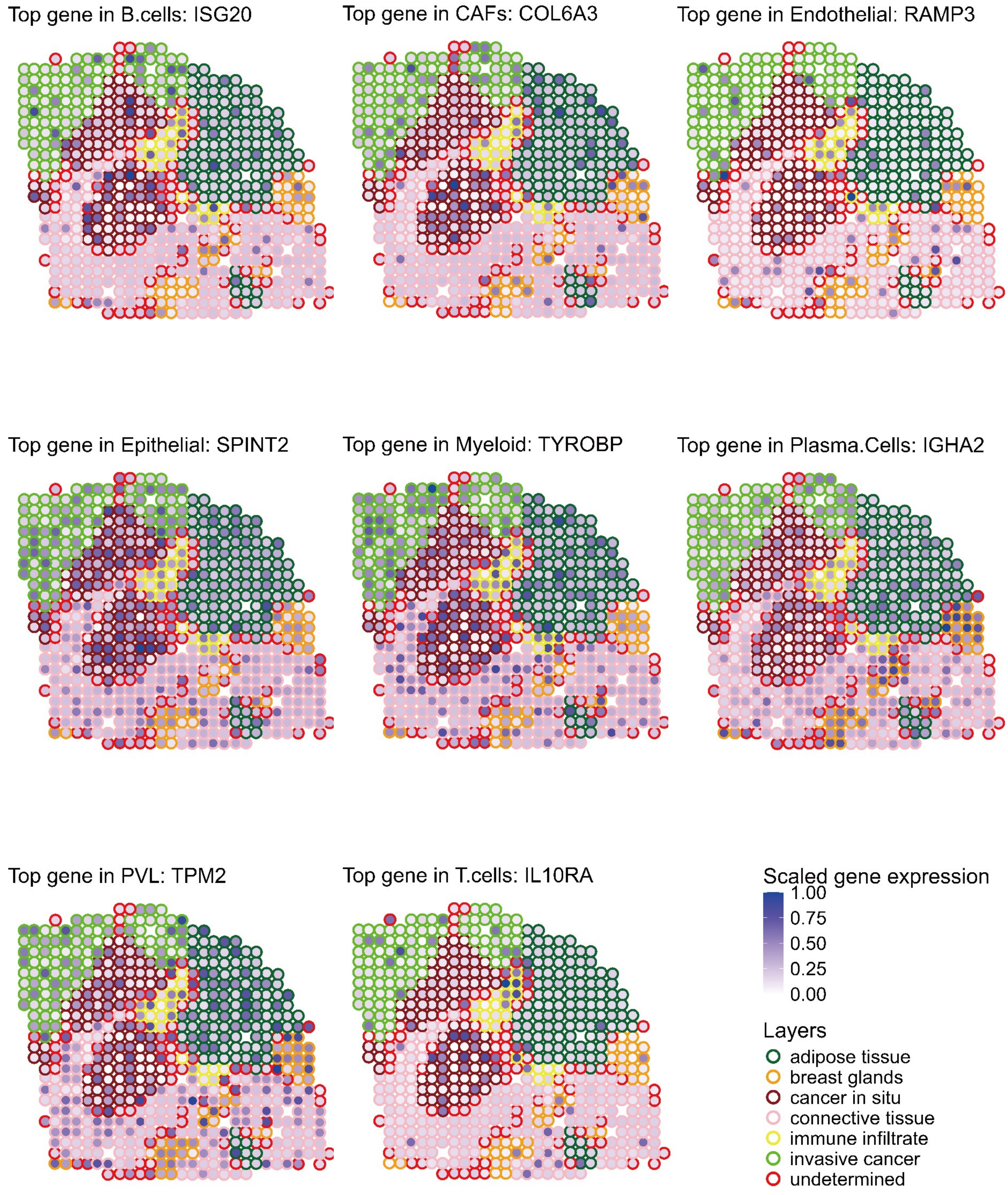


Figure S10: **Spatial expression pattern plots for the top ctSVGs in each cell type for human breast cancer dataset.** Spots are outlined with colors indicating different annotated domains. The scaled gene expression $\tilde{y}_i = \frac{y_i - \min(\mathbf{y})}{\max(\mathbf{y}) - \min(\mathbf{y})}$ is displayed, where $y_i$ is the original gene expression at spot $i$.

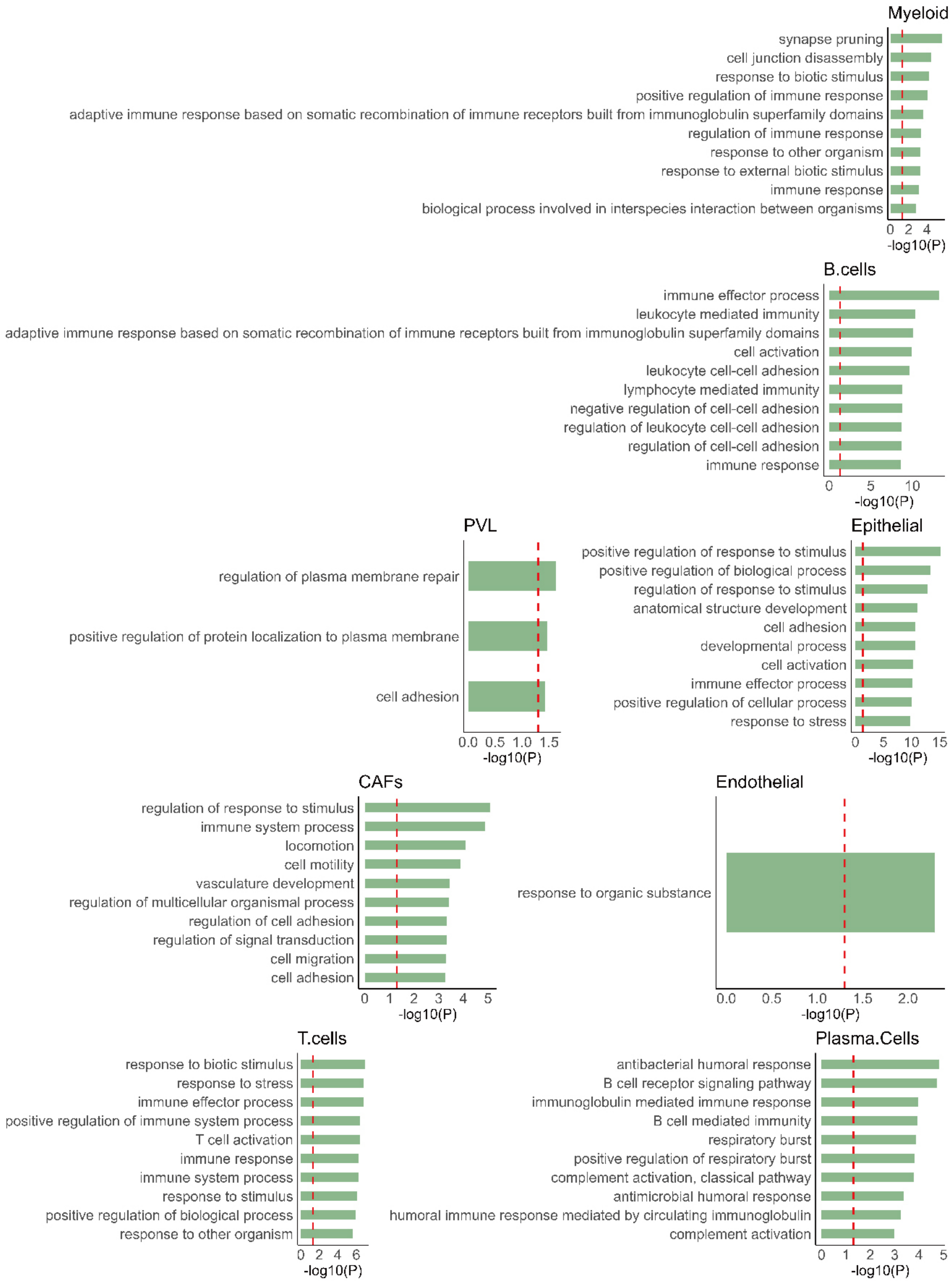


Figure S11: **The gene set enrichment analysis results for the human breast cancer dataset.** The top 10 significant pathways based on ctSVGs detected by STANCE are shown (if the number of significant pathways is less than 10, then all of them are displayed). The enrichment is given as $-\log_{10}$(adjusted p-value), where the default "g_SCS" algorithm is used for multiple testing corrections.

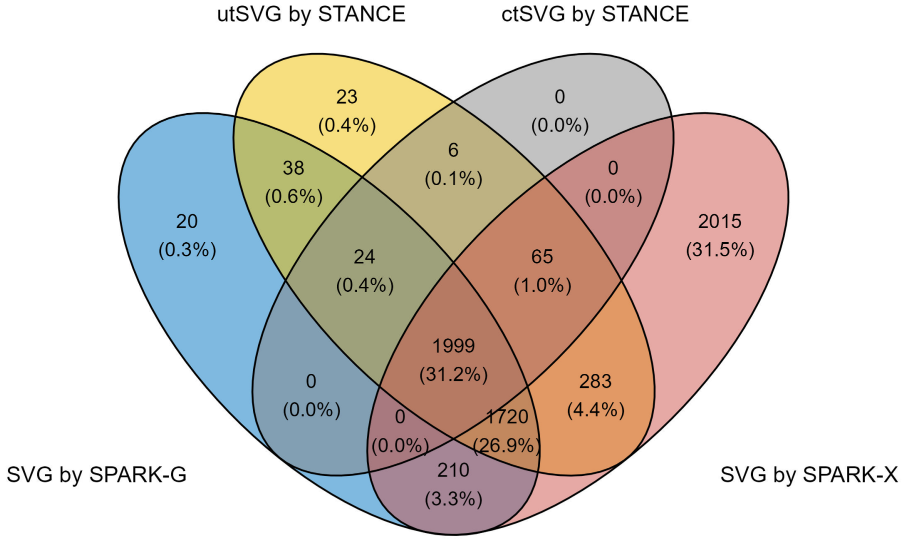


Figure S12: **The Venn diagram for genes identified by different methods in the human kidney cancer dataset.** The Venn diagram shows the logical relationship between sets of genes identified by STANCE, SPARK-G, and SPARK-X. The STANCE overall test identified 4,158 utSVGs (a mixture of SVGs and ctSVGs), with p-values adjusted by the Benjamini-Yekutieli method to meet an FDR rate of 0.01. SPARK-G identified 4,011 SVGs, among which 3,781 were also identified by the STANCE overall test. SPARK-X identified 6,292 SVGs, with 4,067 overlapping with those detected by STANCE. For the 4,158 utSVGs detected by the STANCE overall test, 2,094 ctSVGs were identified across the 12 cell types.

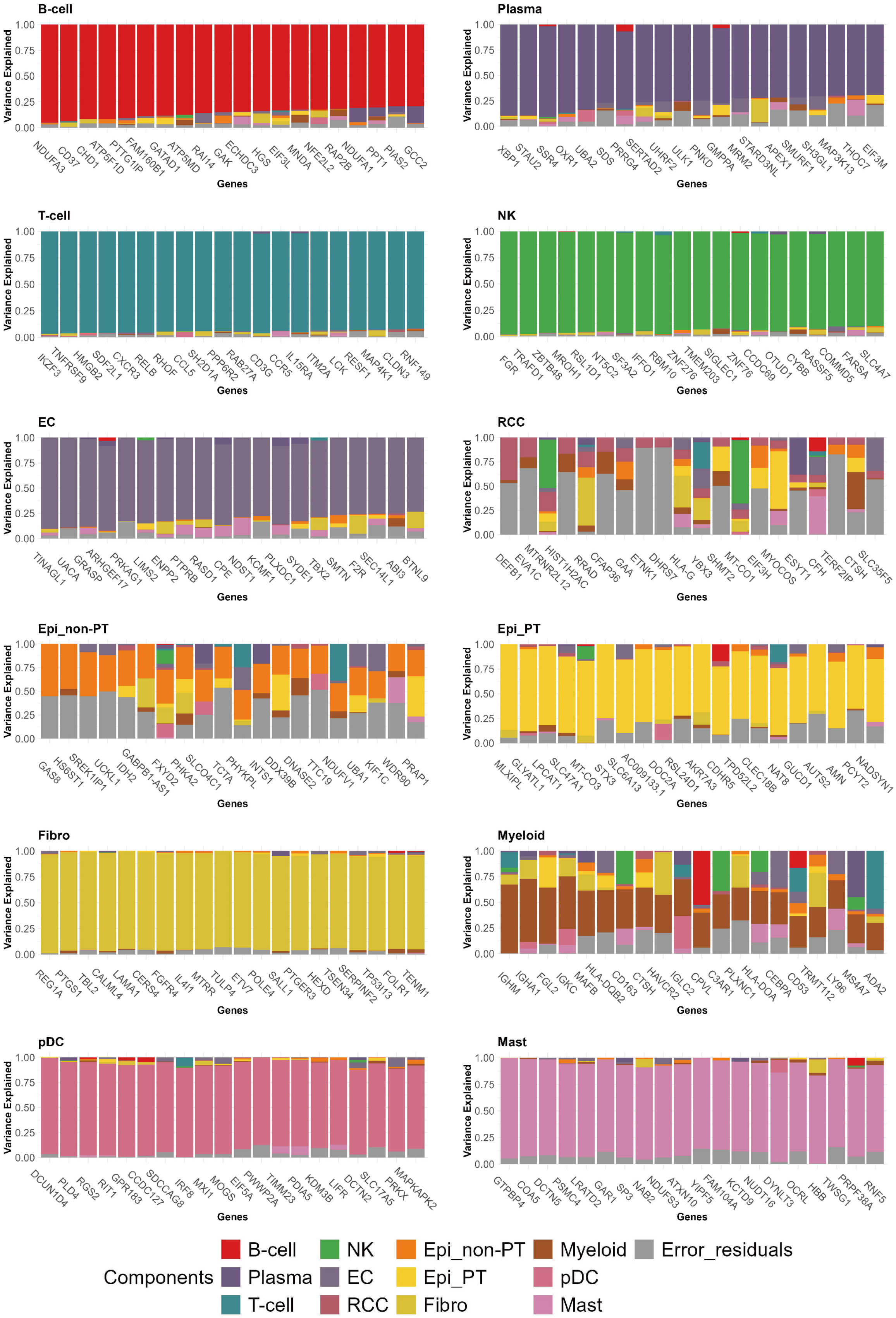
B-cell
Variance Explained
1.00 0.75 0.50 0.25 0.00
NDUFA3 CD37 CHD1 ATP5F1D PTTG1IP FAM160B1 GATAD1 ATP5MD RAI14 GAK ECHDC3 HGS EIF3L MNDA NFE2L2 RAP2B NDUFA1 PPT1 PIAS2 GCC2
Genes
Plasma
Variance Explained
1.00 0.75 0.50 0.25 0.00
XBP1 STAU2 SSR4 OXR1 UBA2 SDS PRRG4 SERTAD2 UHRF2 ULK1 PNKD GMPPA MRM2 STARD3NL APEX1 SMURF1 SH3GL1 MAP3K13 THOC7 EIF3M
Genes
T-cell
Variance Explained
1.00 0.75 0.50 0.25 0.00
IKZF3 TNFRSF9 HMGB2 SDF2L1 CXCR3 RELB RHOF CCL5 SH2D1A PPP6R2 RAB27A CD3G CCR5 IL15RA ITM2A LCK RESF1 MAP4K1 CLDN3 RNF149
Genes
NK
Variance Explained
1.00 0.75 0.50 0.25 0.00
FGR TRAFD1 ZBTB48 MROH1 RSL1D1 NT5C2 SF3A2 IFFO1 RBM10 ZNF276 TMEM203 SIGLEC1 ZNF76 CCDC69 OTUD1 CYBB RASSF5 COMMD5 FARSA SLC4A7
Genes
EC
Variance Explained
1.00 0.75 0.50 0.25 0.00
TINAGL1 UACA GRASP ARHGEF17 PRKAG1 LIMS2 ENPP2 PTPRB RASD1 CPE NDST1 KCMF1 PLXDC1 SYDE1 TBX2 SMTN F2R SEC14L1 ABI3 BTNL9
Genes
RCC
Variance Explained
1.00 0.75 0.50 0.25 0.00
DEFB1 EVA1C MTRNR2L12 HIST1H2AC RRAD CFAP36 GAA ETNK1 DHRS7 HLA-G YBX3 SHMT2 MT-CO1 EIF3H MYOCOS ESYT1 CFH TERF2IP CTSH SLC35F5
Genes
Epi_non-PT
Variance Explained
1.00 0.75 0.50 0.25 0.00
GAS8 HS6ST1 SREK1IP1 UCKL1 IDH2 GABPB1-AS1 FXYD2 PHKA2 SLCO4C1 TCTA PHYKPL INTS1 DDX39B DNASE2 TTC19 NDUFV1 UBA1 KIF1C WDR90 PRAP1
Genes
Epi_PT
Variance Explained
1.00 0.75 0.50 0.25 0.00
MLXIPL GLYATL1 LPCAT1 SLC47A1 MT-CO3 STX3 SLC6A13 AC009133.1 DOC2A RSL24D1 AKR7A3 CDHR5 TPD52L2 CLEC18B NAT8 GUCD1 AUTS2 AMN PCYT2 NADSYN1
Genes
Fibro
Variance Explained
1.00 0.75 0.50 0.25 0.00
REG1A PTGS1 TBL2 CALML4 LAMA1 CERS4 FGFR4 IL4I1 MTRR TULP4 ETV7 POLE4 SALL1 PTGER3 HEXD TSEN34 SERPINF2 TP53I13 FOLR1 TENM1
Genes
Myeloid
Variance Explained
1.00 0.75 0.50 0.25 0.00
IGHM IGHA1 FGL2 IGKC MAFB HLA-DQB2 CD163 CTSH HAVCR2 IGLC2 CPVL C3AR1 PLXNC1 HLA-DOA CEBPA CD53 TRMT112 LY96 MS4A7 ADA2
Genes
pDC
Variance Explained
1.00 0.75 0.50 0.25 0.00
DCUN1D4 PLD4 RGS2 RIT1 GPR183 CCDC127 SDCCAG8 IRF8 MXI1 MOGS EIF5A PWWP2A TIMM23 PDIA5 KDM3B LIFR DCTN2 SLC17A5 PRKX MAPKAPK2
Genes
Mast
Variance Explained
1.00 0.75 0.50 0.25 0.00
GTPBP4 COA5 DCTN5 PSMC4 LRATD2 GAR1 SP3 NAB2 NDUFS3 ATXN10 YIPF5 FAM104A KCTD9 NUDT16 DYNLT3 OCRL HBB TWSG1 PRPF38A RNF5
Genes
Components
B-cell
Plasma
T-cell
NK
EC
RCC
Epi_non-PT
Epi_PT
Fibro
Myeloid
pDC
Mast
Error_residuals

Figure S13: **The stacked variance plots for each cell type in the human kidney cancer dataset.** Displayed are the top 20 significant ctSVGs in each cell type. For each gene, the stacked bar plots display the proportion of variance explained by the 12 cell-type-specific spatial effects and random error. The stacked bar plots indicate that the top 20 ctSVGs of renal cell carcinoma (RCC) cells, non-proximal tubule epithelial (Epi_non-PT) cells and myeloid cells are more heterogeneous in variance than those of the other cell types.

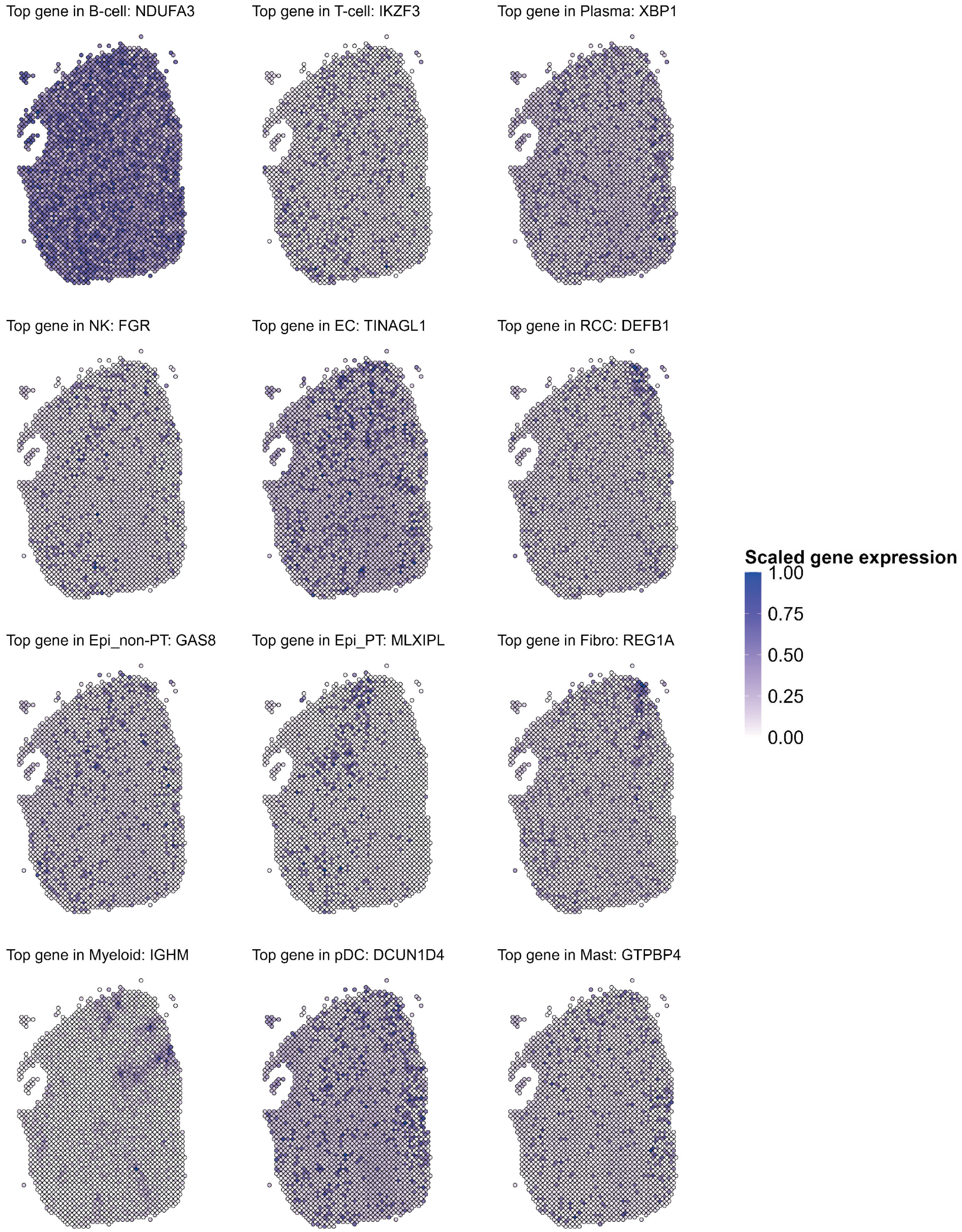


Figure S14: **Spatial pattern plots for the top ctSVGs of each cell type in the human kidney dataset.** The scaled gene expression $\tilde{y}_i = \frac{y_i - \min(\mathbf{y})}{\max(\mathbf{y}) - \min(\mathbf{y})}$ is displayed, where $y_i$ is the original gene expression at spot $i$.

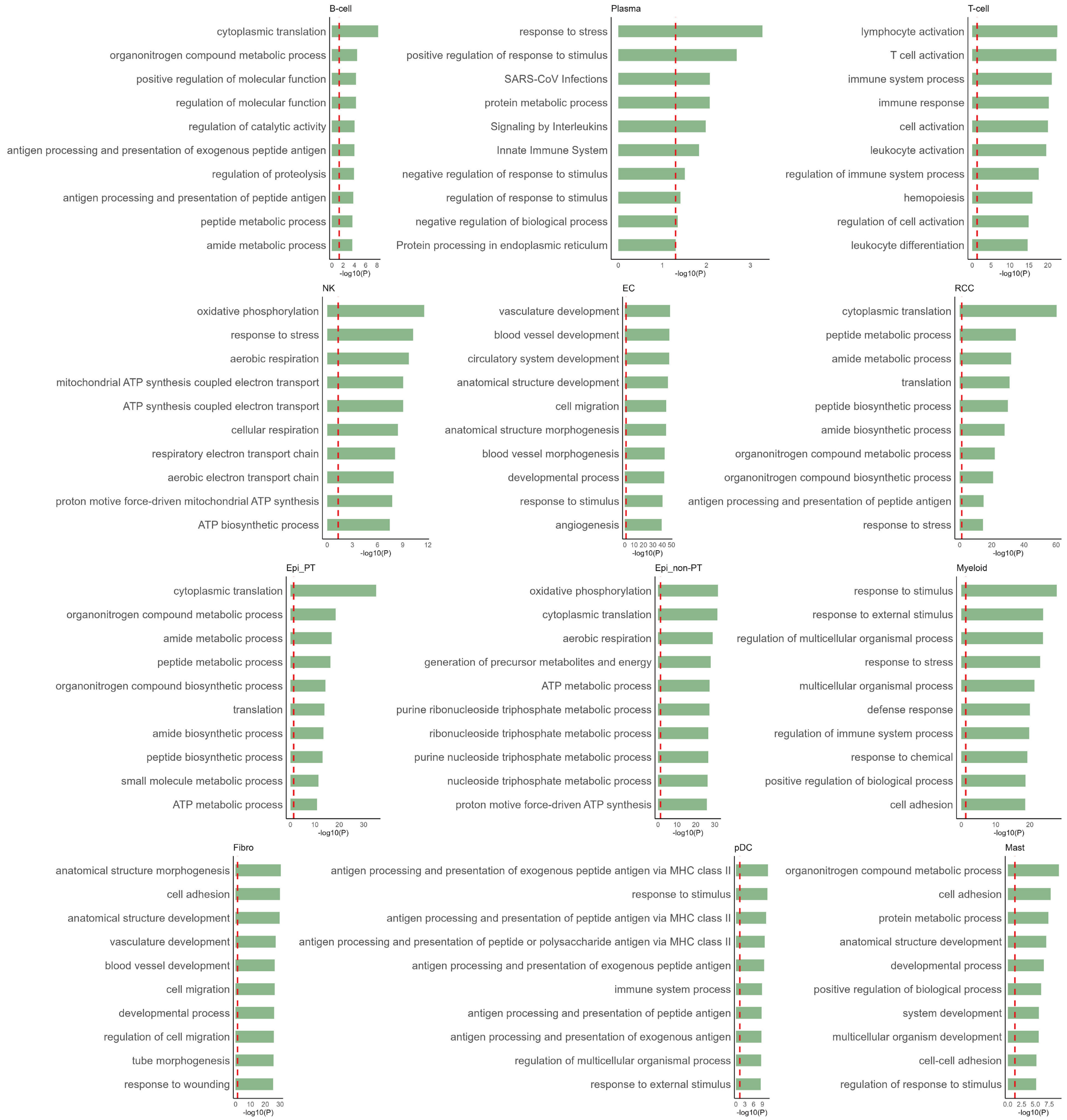


Figure S15: **The gene set enrichment analysis results for the human kidney cancer dataset.** The top 10 significant pathways based on ctSVGs detected by STANCE are shown (if the number of significant pathways is less than 10, then all of them are displayed). The enrichment is given as $-\log_{10}$(adjusted p-value), where the default "g_SCS" algorithm is used for multiple testing corrections.

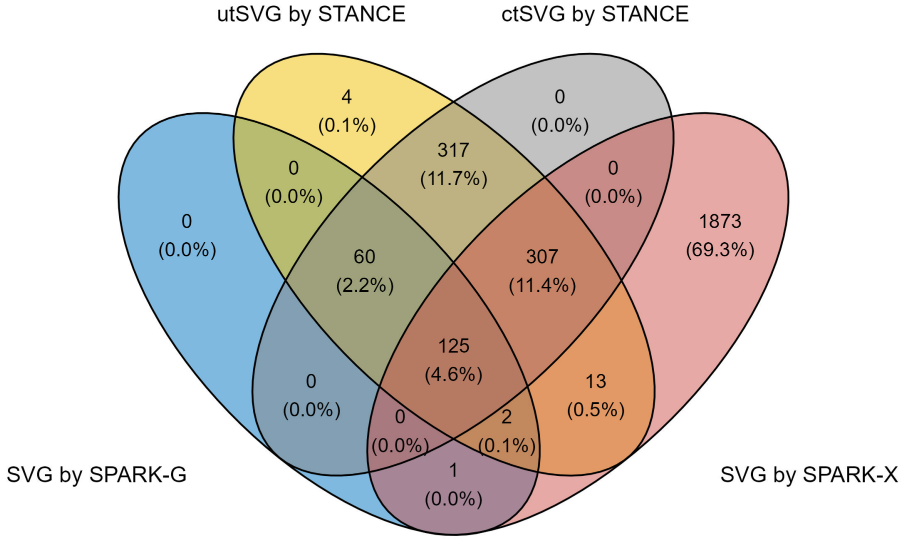


Figure S16: **The Venn diagram for genes identified by different methods in the mouse olfactory bulb dataset.** The Venn diagram shows the logical relationship between sets of genes identified by STANCE, SPARK-G, and SPARK-X. The STANCE overall test identified 828 utSVGs (a mixture of SVGs and ctSVGs), with p-values adjusted by the Benjamini-Yekutieli method to meet an FDR rate of 0.05. SPARK-G identified 188 SVGs, 187 of which were also identified by the STANCE overall test. SPARK-X identified 2,321 SVGs, with 447 overlapping with those detected by STANCE. For the 828 utSVGs detected by the STANCE overall test, 809 ctSVGs were identified across the 12 cell types.

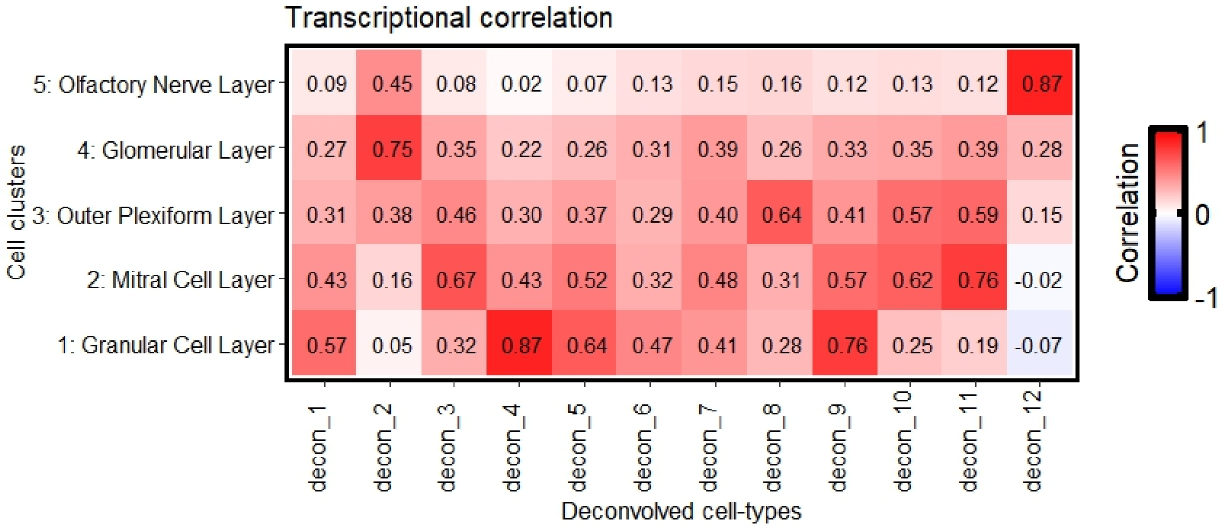


Figure S17: **The transcriptional correlation heatmap for deconvolved cell types and cell clusters in mouse olfactory bulb dataset.** The heatmap visualizes the transcriptional correlation between deconvolved cell types and cell clusters (layers). Specifically, deconvolved cell type 4, cell type 11, cell type 8, cell type 2 and cell type 12 are highly expressed in grangular cell layer, mitral cell layer, outer plexiform layer, glomerular layer and olfactory nerve layer respectively, with highest correlation.

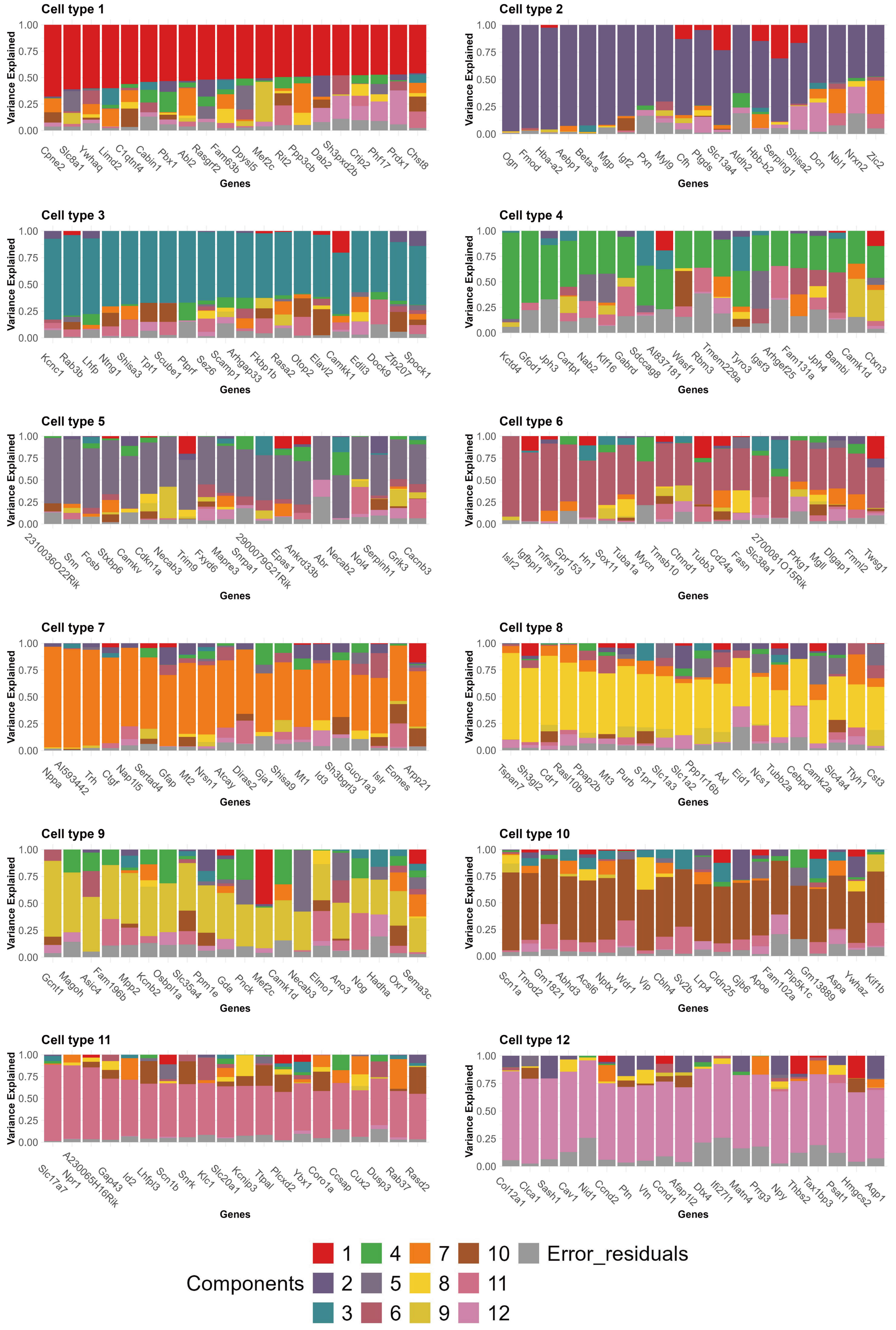
Cell type 1
Variance Explained
1.00 0.75 0.50 0.25 0.00
Cpne2 Slc8a1 Ywhaq Limd2 C1qtnf4 Cabin1 Pbx1 Abl2 Rasgrf2 Fam63b Dpysl5 Mef2c Rit2 Ppp3cb Dab2 Sh3pxd2b Crip2 Phf17 Prdx1 Chst8
Genes
Cell type 2
Variance Explained
1.00 0.75 0.50 0.25 0.00
Ogn Fmod Hba-a2 Aebp1 Beta-s Mgp Igf2 Pxn Myl9 Cfh Ptgds Slc13a4 Aldh2 Hbb-b2 Serping1 Shisa2 Dcn Nbl1 Nrxn2 Zic2
Genes
Cell type 3
Variance Explained
1.00 0.75 0.50 0.25 0.00
Kcnc1 Rab3b Lhfp Ntng1 Shisa3 Tpt1 Scube1 Ptprf Sez6 Scamp1 Arhgap33 Fkbp1b Rasa2 Otop2 Elavl2 Camkk1 Edil3 Dock9 Zfp207 Spock1
Genes
Cell type 4
Variance Explained
1.00 0.75 0.50 0.25 0.00
Kctd4 Gfod1 Jph3 Cartpt Nab2 Kif16 Gabrd Sdccag8 AI837181 Wasf1 Rbm3 Tmem229a Tyro3 Igsf3 Arhgef25 Fam131a Jph4 Bambi Camk1d Ctxn3
Genes
Cell type 5
Variance Explained
1.00 0.75 0.50 0.25 0.00
2310036O22Rik Snn Fosb Stxbp6 Camkv Cdkn1a Necab3 Trim9 Fxyd6 Mapre3 Snrpa1 2900079G21Rik Epas1 Ankrd33b Abr Necab2 Nol4 Serpinh1 Grik3 Cacnb3
Genes
Cell type 6
Variance Explained
1.00 0.75 0.50 0.25 0.00
Islr2 Igfbpl1 Tnfrsf19 Gpr153 Hn1 Sox11 Tuba1a Mycn Tmsb10 Ctnnd1 Tubb3 Cd24a Fasn Slc38a1 2700081O15Rik Prkg1 Mgll Dlgap1 Fmnl2 Twsg1
Genes
Cell type 7
Variance Explained
1.00 0.75 0.50 0.25 0.00
Nppa AI593442 Trh Ctgf Nap1l5 Sertad4 Gfap Mt2 Nrsn1 Atcay Diras2 Gja1 Shisa9 Mt1 Id3 Sh3bgrl3 Gucy1a3 Islr Eomes Arpp21
Genes
Cell type 8
Variance Explained
1.00 0.75 0.50 0.25 0.00
Tspan7 Sh3gl2 Cdr1 Rasl10b Ppap2b Mt3 Purb S1pr1 Slc1a3 Slc1a2 Ppp1r16b Axl Eid1 Ncs1 Tubb2a Cebpd Camk2a Slc4a4 Ttyh1 Cst3
Genes
Cell type 9
Variance Explained
1.00 0.75 0.50 0.25 0.00
Gcnt1 Magoh Asic4 Fam196b Mpp2 Kcnb2 Osbpl1a Slc35a4 Ppm1e Gda Pnck Mef2c Camk1d Necab3 Elmo1 Ano3 Nog Hadha Oxr1 Sema3c
Genes
Cell type 10
Variance Explained
1.00 0.75 0.50 0.25 0.00
Scn1a Tmod2 Gm1821 Abhd3 Acsl6 Nptx1 Wdr1 Vip Cbln4 Sv2b Lrp4 Cldn25 Gjb6 Apoe Fam102a Pip5k1c Gm13889 Aspa Ywhaz Kif1b
Genes
Cell type 11
Variance Explained
1.00 0.75 0.50 0.25 0.00
Slc17a7 Npr1 A230065H16Rik Gap43 Id2 Lhfpl3 Scn1b Snrk Klc1 Slc20a1 Kcnip3 Tlpal Plcxd2 Ybx1 Coro1a Ccsap Cux2 Dusp3 Rab37 Rasd2
Genes
Cell type 12
Variance Explained
1.00 0.75 0.50 0.25 0.00
Col12a1 Clca1 Sash1 Cav1 Nid1 Ccnd2 Ptn Vtn Ccnd1 Afap1l2 Dtx4 Ifi27l1 Matn4 Prg3 Npy Thbs2 Tax1bp3 Psat1 Hmgcs2 Aqp1
Genes
Components
1 2 3 4 5 6 7 8 9 10 11 12 Error_residuals

Figure S18: **The stacked variance plots for each cell type in the mouse olfactory bulb dataset.** Displayed are the top 20 significant ctSVGs in each cell type. For each gene, the stacked bar plots display the proportion of variance explained by the 12 cell-type-specific spatial effects and random error.

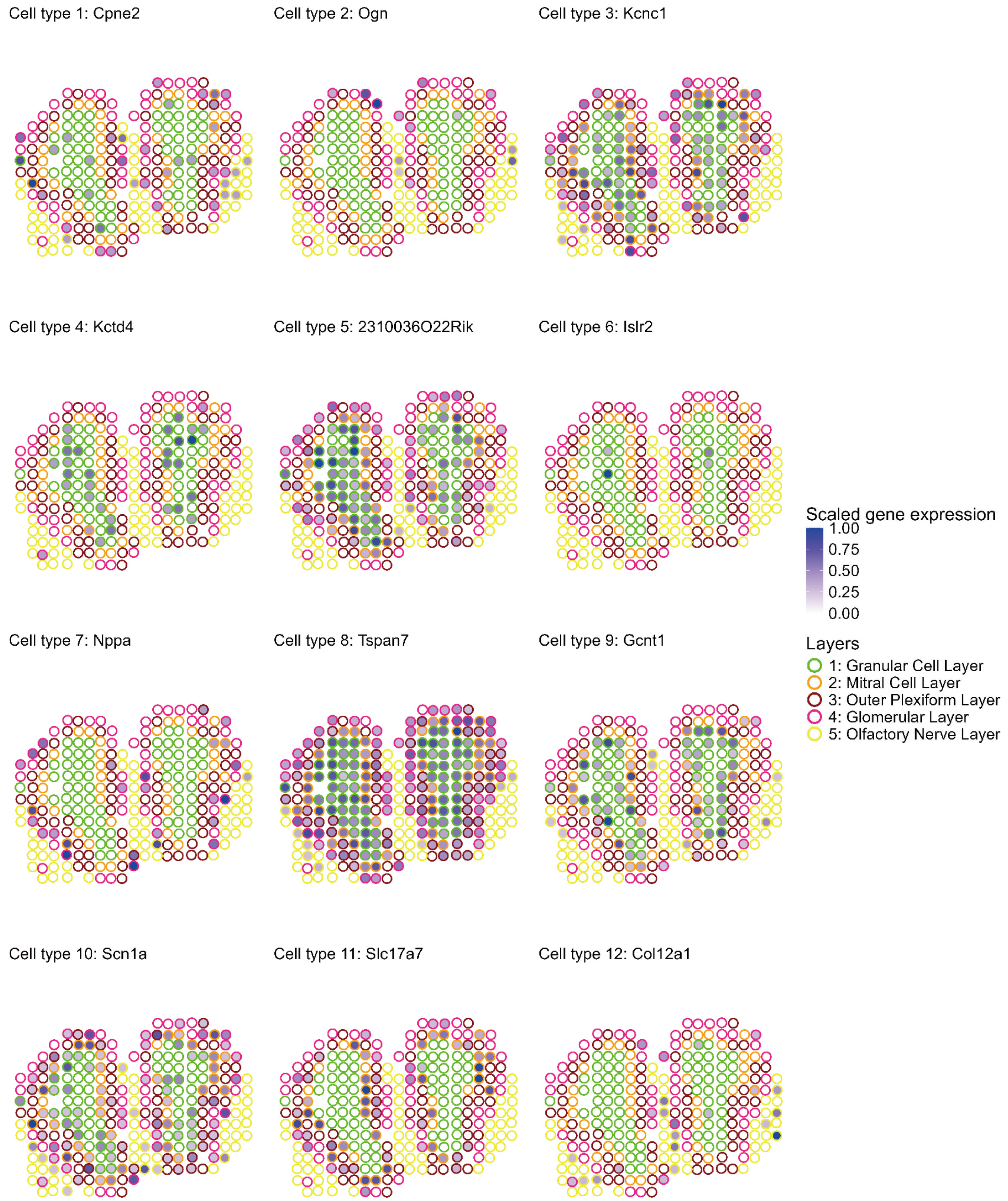


Figure S19: **Spatial pattern plots for the top ctSVGs of each cell type in the mouse olfactory bulb dataset.** Spots are outlined with colors indicating the annotated domains. The scaled gene expression $\tilde{y}_i = \frac{y_i - \min(\mathbf{y})}{\max(\mathbf{y}) - \min(\mathbf{y})}$ is displayed, where $y_i$ is the original gene expression at spot $i$.
